# Improving anxiety research: novel approach to reveal trait anxiety through summary measures of multiple states

**DOI:** 10.1101/2023.06.01.543235

**Authors:** Zoltán K Varga, Diána Pejtsik, Máté Tóth, Zoltán Balogh, Manó Aliczki, László Szente, Gyula Y Balla, Levente Kontra, Zsófia Eckert, Huba Szebik, Zsolt Borhegyi, Éva Mikics

## Abstract

The reliability and validity of preclinical anxiety testing is essential for translating animal research into clinical use. However, the commonly used anxiety tests lack inter-test correlations and face challenges with repeatability. While translational animal research should be able to capture stable individual anxiety traits, the current approach employs a single type of test at a single time that only measures transient states of animals, heavily influenced by experimental conditions. Here, we propose a validated, optimized test battery capable of reliably capturing trait anxiety in rats and mice of both sexes. Instead of developing novel tests, we combined widely used tests (elevated plus-maze, open field and light-dark test) to provide instantly applicable adjustments for better predictive validity. We repeated these tests three times to capture behaviour across multiple challenges, which we combined to generate summary measures (SuMs). Our approach resolved between-test correlation issues and provided better predictions for subsequent outcomes under anxiogenic conditions or fear conditioning. SuMs were also able to reveal differences in anxiety in an etiological stress model. Finally, we tested our method’s efficacy in discovering anxiety-related molecular pathways through RNA sequencing of the medial prefrontal cortex. Using SuMs, we identified four-times more molecular correlates of trait anxiety as compared to transient anxiety states, pointing out novel functional gene clusters. Furthermore, 16% of these molecular correlates of anxiety were replicated in amygdalar samples, as well. In summary, we provide a novel approach to capture trait anxiety in rodents, offering improved predictions for potential therapeutic targets for personalized medicine.

## Introduction

Most of us have experienced anxiety during our lives. The tendency to have such incidents stems from our individually characteristic, stable trait anxiety (TA)^1^. High levels of TA predict a high frequency of anxious states (SA) in different situations and are the determining core of anxiety disorders^2,3^. As the common root of the autonomic, emotional and behavioral manifestation of transient SAs, TA represents the optimal psychiatric intervention target^4^.

In pathological conditions of anxiety, anxiolytic therapy is ineffective in 4 out of 10 clinical cases, leaving anxiety disorders the most prevalent and burdening mental illnesses^5^. Prescribed compounds act through brain mechanisms that have been targeted for at least 30 years with moderate success, and new candidates are rare^6^. Unfortunately, animal models, our primary source of information on the molecular basis of anxiety, offer targets that fail in clinical trials in more than 60% of the cases^7^. Such alarming rates of failed medicine candidates at clinical and preclinical levels suggest that our approach to investigating anxiety is in a crisis and needs fundamental improvement.

This translational gap potentially stems from the fact that anti-anxiety therapies are considered satisfactory if they control the permanent TA, however, current animal tests only detect the fluctuating SA^4,8,9^. Consequently, we can only expect these tests to highlight markers and treatment targets of transient symptoms rather than drivers of the underlying phenomenon. Results from the most commonly used^7^ rodent paradigms, the elevated plus-maze (EPM)^10,11^, the light-dark (LD)^12,13^ and the open field (OF)^14,15^ tests, support this idea. Behavioral parameters measured in these tests are not correlated with each other^4,16^ and are hardly reproducible^17–19^, which indicates a lack of a shared basis for these, such as TA. Alternative approaches investigate TA using selectively bred highly anxious animal strains^20–22^. Unfortunately, these approaches i) are only suitable for the examination of TA of that specific strain, ii) their molecular profile is not necessarily in connection with the anxious phenotype and iii) are based on the low-validity tests mentioned above. Consequently, despite the alarming flaws of current methods, and the urgent need for a proper one, the field lacks an adequate tool to measure the most relevant subject of anxiety research: TA.

To bridge this gap, we developed an approach to measure TA that is based on refined sampling and analysis of conventional anxiety tests. We hypothesize that the behavior we measure in these tests is the direct consequence of SAs, which emerge as the interaction of the individually stable TA and the current environmental context (Fig.1 – hypothesis). Assuming this, improved sampling of SAs converge to TA. Our approach uses the conventional anxiety tests but expands the classical sampling method in time and depth to represent multiple states of our subjects (Fig.1 – sampling design) by measuring i) more variables, ii) involving serial testing events and even iii) more test types in our analysis. Particularly, we sampled SA in rodents with different sequences of the most frequently used anxiety tests, where each test was repeated three times with all animals. We measured a range of variables (single measures, SiMs), assessed the possibility of their summarisation, and created summary measures (SuMs) by averaging SiMs of serial tests and composite measures (COMPs) by averaging SiMs of different test types (Fig.1 – analysis design). In medical research, creating SuMs of serial sampling is one effective method for describing the background construct of fluctuating measures while decreasing environmental noise and avoiding over-parametrisation^23^. Despite the advantages of such an approach, the field of behavioral neuroscience almost entirely misses using it. We validated SuMs as measures of TA with multiple novel approaches. i) Since TA is assumed to be the shared basis of different anxiety tests, we hypothesized that SuM-based inter-test correlations would be stronger than the classical SiM-based ones. ii) Since TA is stable across time and contexts, we hypothesized that SuMs better predict acute stress-induced future responses, such as behavior in anxiety or fear-triggering environments. iii) Since high levels of TA is a core symptom of anxiety disorders, we hypothesized that SuMs are more sensitive markers of chronic stress-induced anxiety in an etiological stress model. Following these validation steps, we examined the sensitivity of SuMs in discovering molecular markers and potential therapeutic candidates for anxiety by conducting an RNAseq experiment in the anxiety-mediator area medial prefrontal cortex (mPFC) of resting state animals.

**Figure 1.**
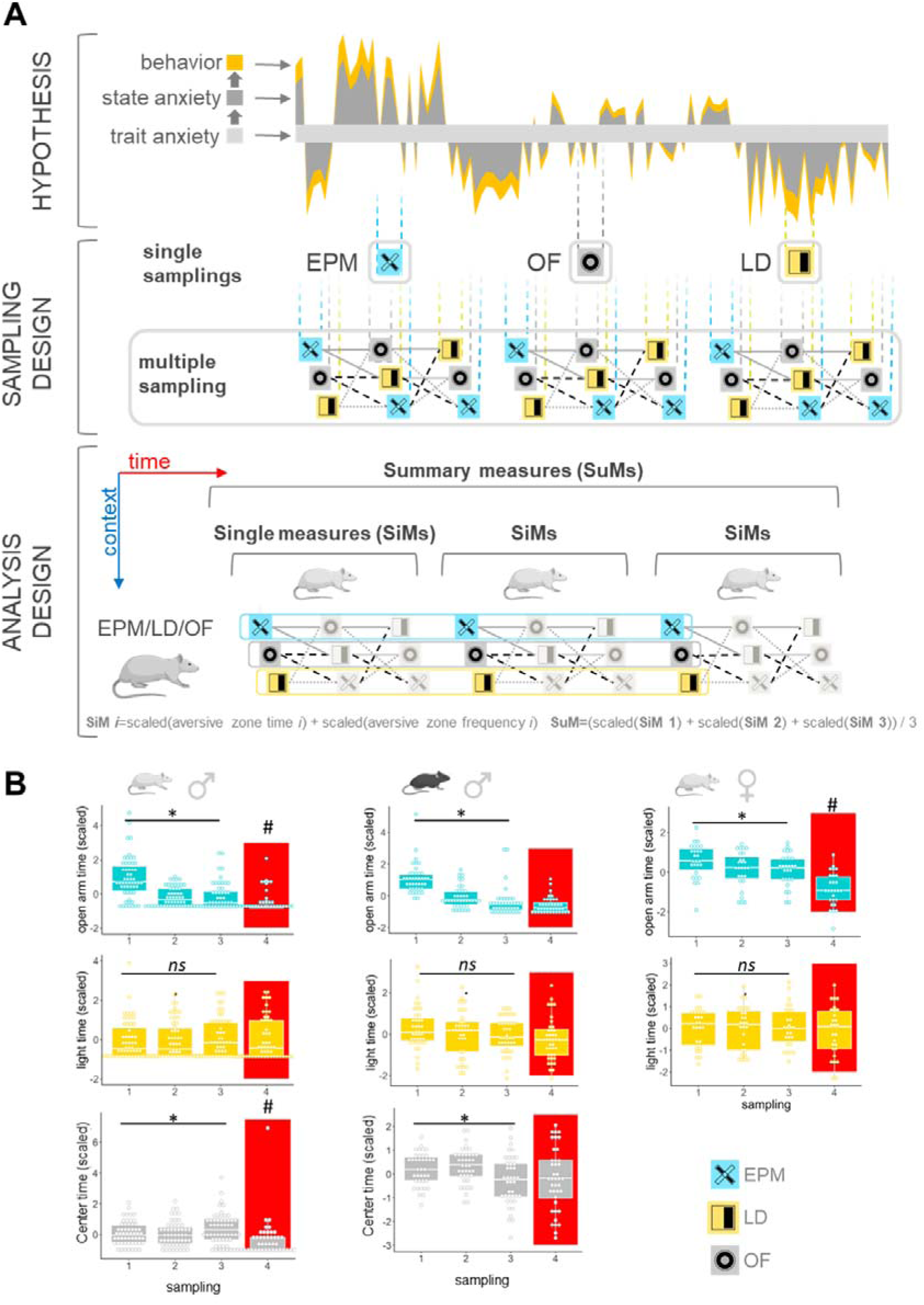
**A) Hypothesis, sampling, and analysis design. Hypothesis:** Schematic drawing shows how an individual’s behavior is expressed as a consequence of state and trait anxiety according to our working hypothesis. Measured anxiety-like behavior is an indirect consequence of trait anxiety and is influenced by time and context, hence using a single anxiety test leads to undersampled experiments. **Sampling design:** we used repeated sampling with multiple test-types in a semi-random order to cover more internal and external variability stemming from the individual’s trait, state, environmental context, and their fluctuation in time. **Analysis design:** Behavioral data (time spent and frequency of enters to the aversive compartment) from each test were used to create classical single measures (SiM) and more stable summary measures (SuM), by combining multiple variables or testing events to describe trait anxiety. B) **Shared and unique features of anxiety tests in rats.** The effect of test repetitions on the most frequently used anxiety-like behavioral outcome (time spent in the aversive compartment) of the EPM, LD, and OF anxiety tests in male (left) and female (right) rats and male mice (middle). Red rectangles highlight the testing event in aversive conditions. All statistical results are presented in Table 1. **Abbreviations:** EPM: elevated plus-maze test, OF: open field test, LD: light-dark test * p<0.05 test repetition main effect. #: p<0.05 significant difference between the aversive and the last baseline sampling event

**Table 1.:**
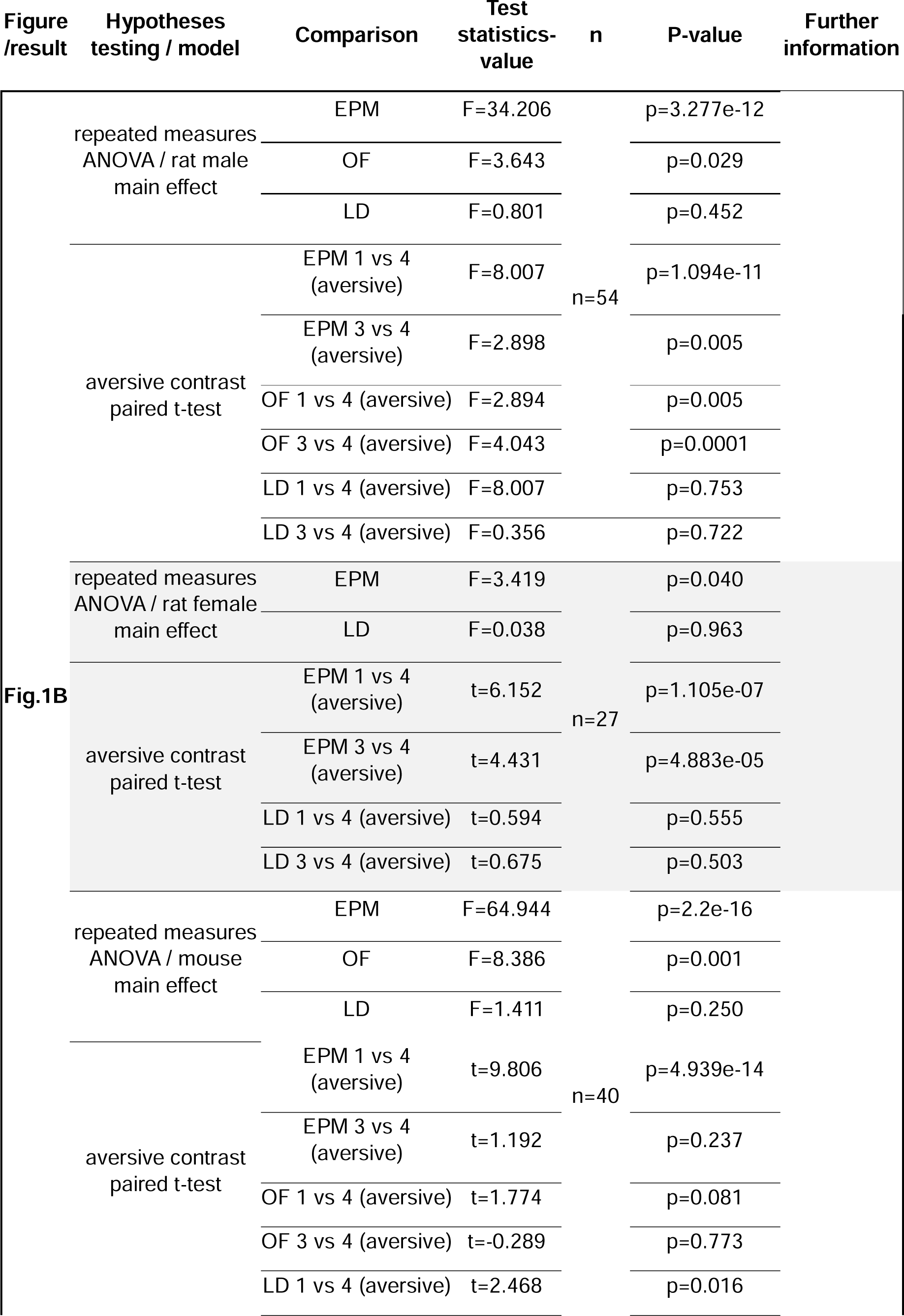
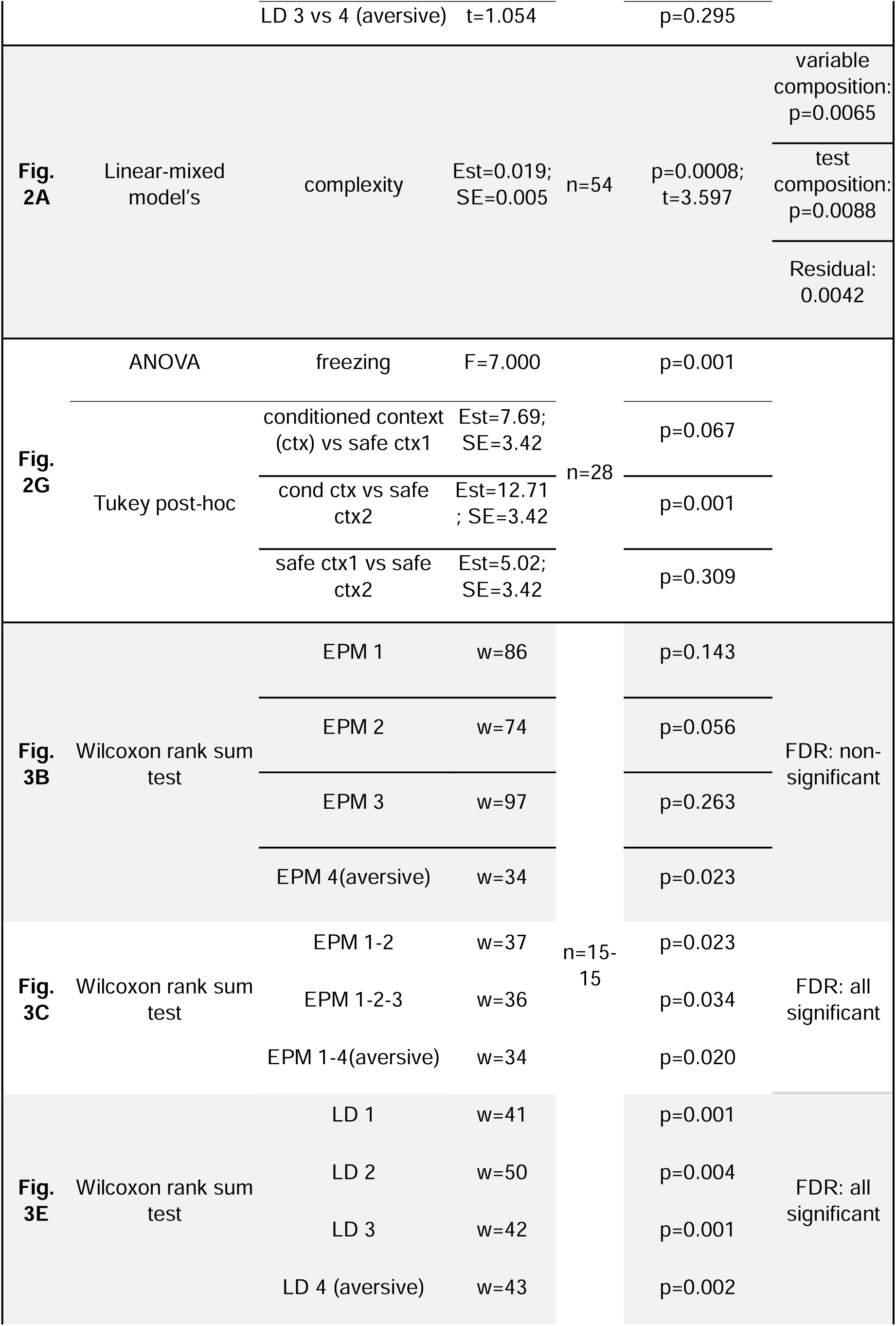
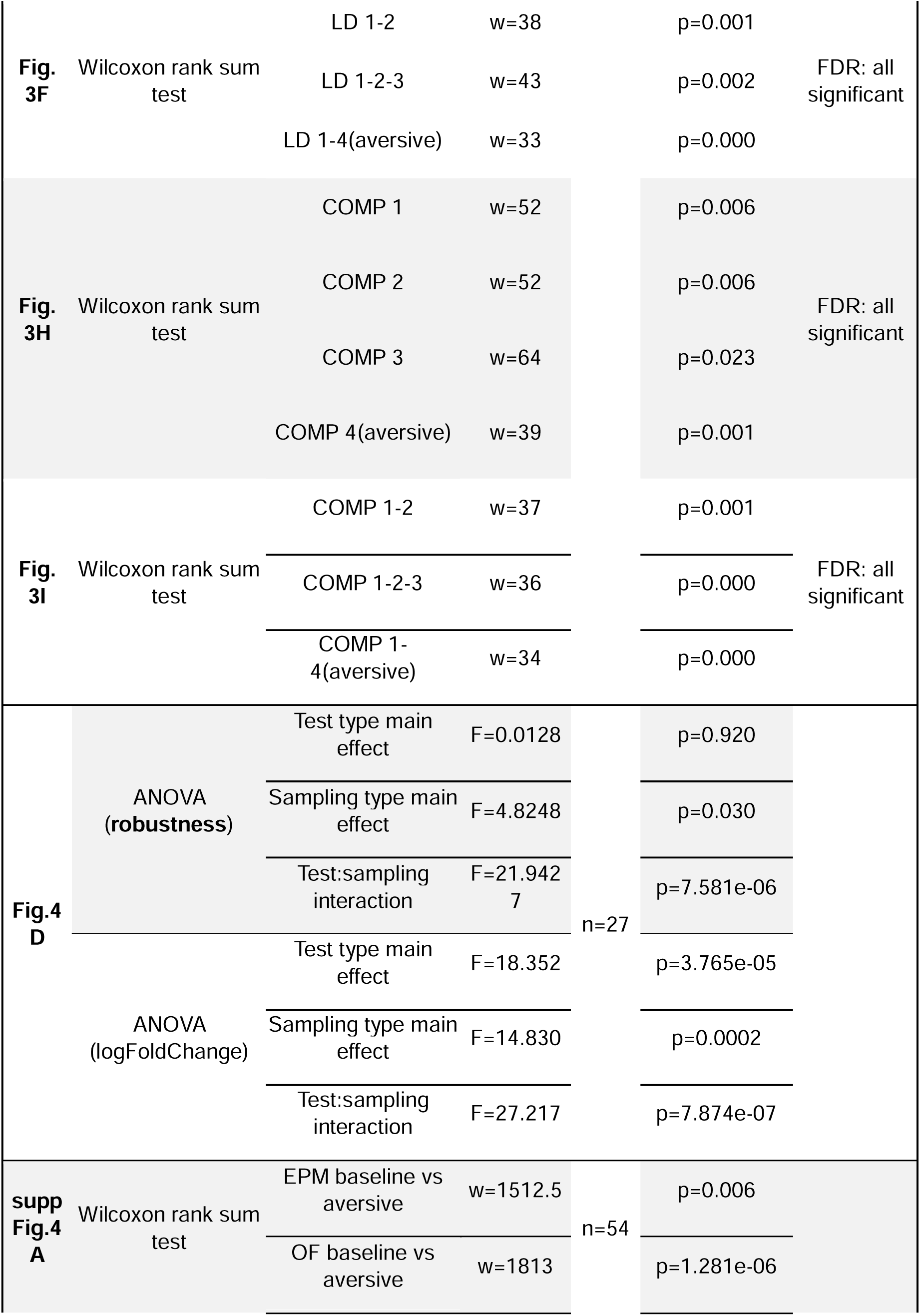
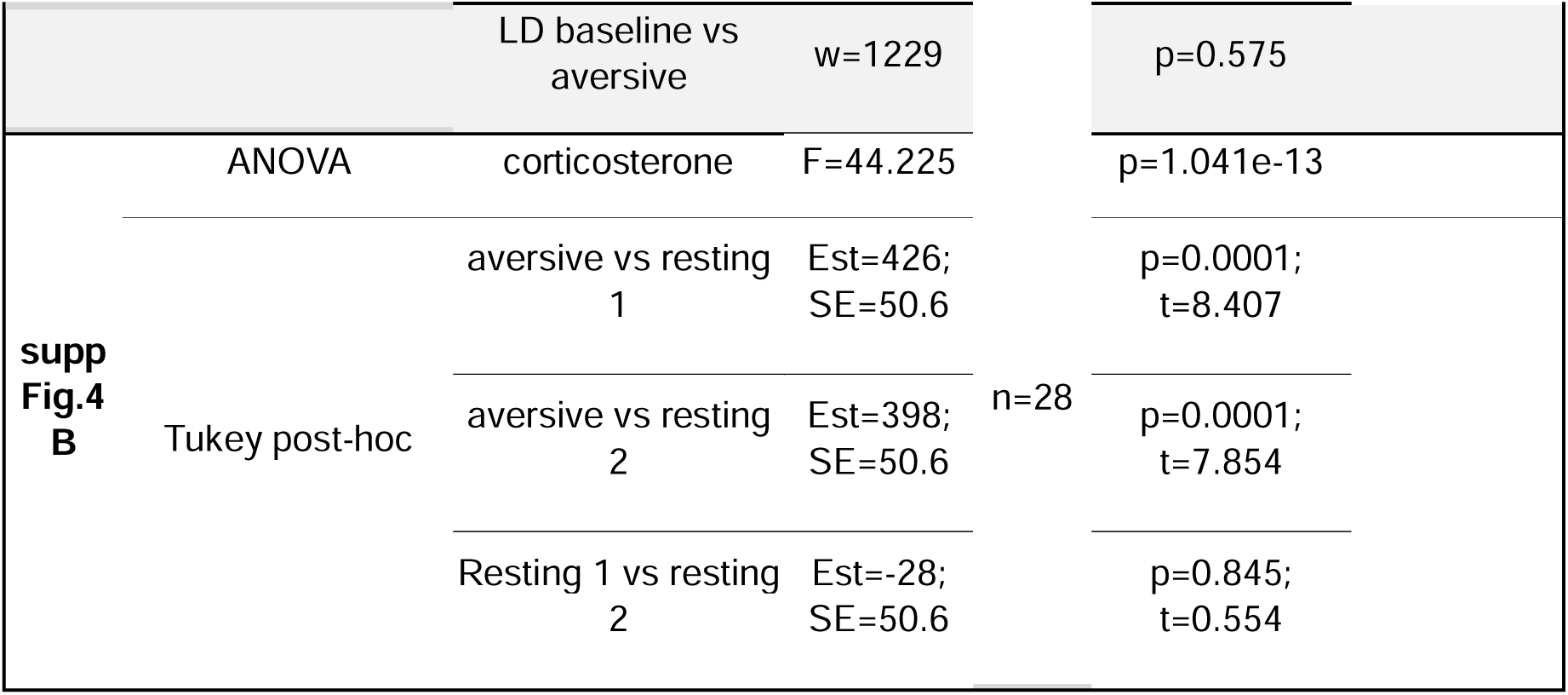
Parameters of test statistics. Table 1 shows (left-to-right) the result section/figure number corresponding to the statistical parameters, the hypotheses testing method/model that were used, the comparisons that were made, and the resulting test statistics-values, sample sizes, and significance levels. We presented the conventional parameters of the particular statistical approaches such as F values in the case of ANOVAs, t or w values in the case of t-tests or Wilcoxon tests, as well as estimates (est) and standard errors (SE) in the case of linear mixed models. The column entitled Further information shows additional relevant information that is specific for a particular method, like random variances in mixed models or the results of FDR correction following multiple comparisons.

## Results

### Overlaps between anxiety-like variables support the use of SuMs

If variables carry partially overlapping information, summarisation is possible and also beneficial for describing their shared background. We assessed the level of overlap between EPM, LD, and OF test variables by repeatability (RA) and correlation analysis in rats and mice. We described repeatability with test-retest changes in baseline and more aversive lighting conditions, within and between-test variability, and bootstrap analysis-based repeatability measures. Test-retest changes were evident in the EPM and the OF, but not in the LD test (Fig.1B). Similarly, aversivity increased anxiety in the EPM and OF, but not in the LD test. Between-test variance were smaller in OF and LD while within-test variance were the highest in LD variables indicating robustness and sensitivity for individual differences. Repeatability measures confirmed such ranking of the tests (supp.Fig.1A,2B). Variables of the same test type clustered together in repeatability. However, frequency variables were the most, while latency variables were the least reproducible variables in most cases, indicating a similarity between the tests. PCA revealed further conformities, as variables of all tests were loaded in the same dimension (supp.Fig.1B,2C). These trends were similar in both species and sexes. In summary, there are test type and variable type similarities of these variables which are consistent in the investigated species and sexes. This indicates a shared but differently expressed basis of these anxiety tests and potential in the summarisation of their variables.

### SuMs reveal shared variance of anxiety tests and predict acute stress-induced behaviour

Since variables of the same anxiety test were similar in PCA and RA, we averaged these from multiple testing events into SuMs to describe TA. As TA is the permanent basis of anxiety-like responses^2,3^, TA markers should be expressed consistently through contexts and time. To test this, we created differentially complex SuMs of the EPM, LD, and OF tests from a series of repeated design experiments. We assessed i) their inter-test correlations and their efficacy to predict behaviour in different stress-inducing contexts such as ii) the same tests in an aversive condition (Fig.2D-E), iii) fear conditioning (CFP)(Fig.2F-H), or iv) acoustic startle paradigms (Fig.2H). Our simplest variables (SiMs) were the time spent in, enter frequency or latency to the aversive compartment of a test sampled ones. Our most complex SuMs consist of all of these variables sampled three times. We constructed all possible variables along this scale and described those with a complexity score (=number of variables x number of testing events).

**Figure 2.**
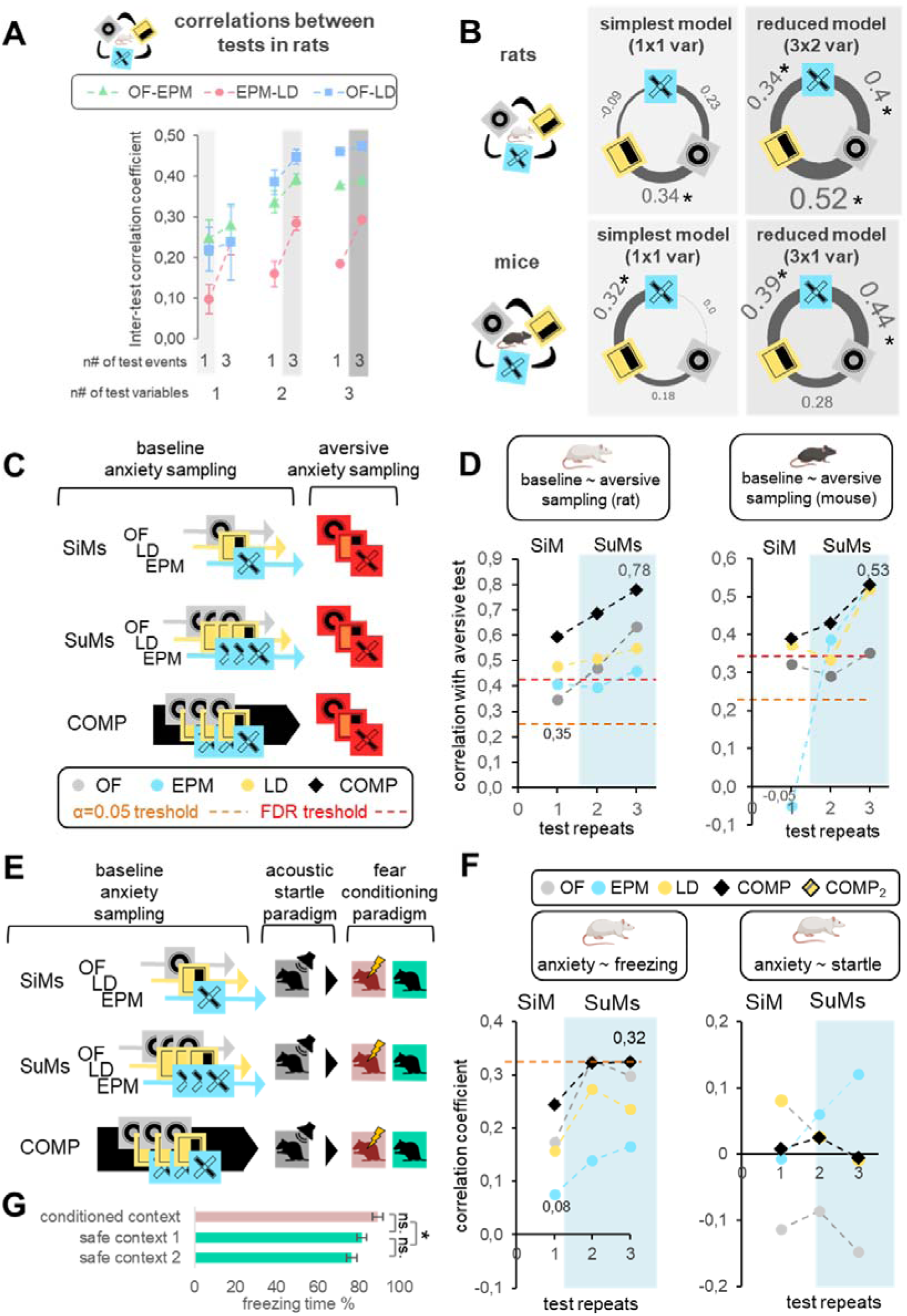
Inter-test correlations and behavioral predictions as a function of variable complexity. **A)** Inter-test correlations between increasingly complex variables of different anxiety tests. The correlation coefficients between OF and EPM (green), EPM and LD (red), and LD and OF (blue) were plotted at different variable combinations of these tests. The #n of test events and the #n of test variables refers to how many repeated test events (1 or 3 repeats) were included and to how many different variable types (time, frequency, latency) were used to create summary variables, respectively. SiMs are single variables, for example time spent in the open arms of an EPM, while SuMs consist of either multiple samplings of one variable type and/or multiple variable types measured in one or multiple events/repeats. **B)** Network plot of inter-test correlations between different tests revealed by SiMs (simplest model) and SuMs (reduced) in the rat (upper) and mouse (lower) experiments. **C) Behavioral predictions of anxiety measures.** Correlation coefficients were measured between identically composed variables in baseline and aversive contexts. SiMs represent one, and SuMs represent more testing events, while COMPs include SiMs or SuMs of all test types. **D) Correlations between increasingly complex anxiety measures** (SiMs, SuMs, or COMPs) from the baseline sampling and their aversive counterparts in rats (left) and mice (right). **E)** Correlation coefficients were measured between baseline anxiety measures (SiMs, SuMs, COMPs), freezing in the fear conditioning paradigm, and startle in the acoustic startle paradigm. **F)** Time spent with freezing (%) in the conditioned fear paradigm in the contextual reminder (conditioned) and safe (different) contexts. Freezing was sampled two times in the safe context, from which an average freezing value was used to be predicted. **G) Correlation of increasingly complex anxiety measures** (SiMs, SuMs, or COMPs) with freezing behavior in novel context following fear conditioning (left) and startle response to acoustic stimuli (right). COMP represents the composite of all anxiety tests and measures. Red dashed lines represent the threshold of significance on the scale of correlation coefficients. Additional numbers on the plots show the smallest and largest coefficients. All statistical results are presented in Figure 5A). Fig 2G) can be found in Table 1.

We found that SuMs of high complexity correlated stronger between tests than less complex SuMs or the SiMs in both rats and mice (Fig.2A-C)(Fig.5A). Inter-test correlations could be explained by variable complexity as a fixed effect, and variable and test composition as interacting random effects. The weakest correlations come from the least complex SuMs, and SuMs that include latencies, while the strongest ones consist of more complex and/or frequency and time-included data (suppFig.3). We excluded latencies from further analysis and used reduced models with time and frequency data which also outperformed the SiM variables. The association between variable complexity and inter-test correlations were apparent in both species (Fig.2A-C, Fig.5A).

In aversive sampling, we detected enhanced anxiety-like behavior (Fig.2D-E) and corticosterone response compared to baseline conditions (Supplementary fig.4B). Correlation analysis between test-specific SiMs and SuMs versus their aversive counterparts revealed that more complex variables predict future behavioral responses better in both species (Fig.2E). Furthermore, creating composite scores from SiMs and SuMs of all tests (COMP) outperformed test-specific SiMs and SuMs, respectively, reaching a 0.78 correlation between baseline and aversive responses. Predictions of SuMs were higher than the predictions of any of our SiMs (1^st^, 2^nd,^ or 3^rd^ testing) indicating that their performance is not due to a carry-over effect of repeated testing but these variables explain additional information. Likewise, the predictions of COMPs were higher than the predictions of SuMs which indicates that those carry additional information to any test type that we used here. Similar to our previous studies, baseline or stress-induced corticosterone levels were not correlated with anxiety variables^38^. We confirmed these findings in both species and sexes, showing a similar association between the number of sampling events and their predictive power (Fig.5A).

We also aimed to predict the amount of freezing behavior in a safe context following CFP as a measure of generalized fear, a symptom of post-traumatic stress disorder^39^. Anxiety tests showed poor performance in predicting fear, but we observed a similar increase in correlations in response to using more complex variables, especially COMPs (Fig.2H)(Fig.5A). Acoustic startle responses could not be predicted by either variable, regardless of the sampling approach. These results indicate that SuMs and their composites can predict specific future states of the animals entering challenging environments.

### SuMs are more sensitive markers of chronic stress-induced behaviour

Since TA is the core symptom of anxiety disorders, TA markers should be responsive to etiological models of such conditions. Post-weaning social isolation (PWSI) is an extensively used animal model of childhood neglect, a prevalent maltreatment that facilitates anxiety^40^. To evaluate the sensitivity of our approach in characterising isolation-induced anxiety, we reduced our protocol to three baselines and one aversive sampling with the EPM and LD tests following PWSI. In EPM, we found no isolation-related differences on any sampling days (Fig.3B). However, SuMs were higher following PWSI: two tests were necessary to show isolation-induced anxiety (Fig.3C). Since all LD and COMP variables were significantly increased by isolation (Fig.3E,3F,3H,3I), we also compared its effect sizes on these measures. We found that the effect sizes based on averaged sampling events (SuMs) were higher than the averages of effect sizes of those very same events. In other words, SuMs were more sensitive to isolation-induced anxiety than any SiMs (Fig.3D,3G,3J).

**Figure 3.:**
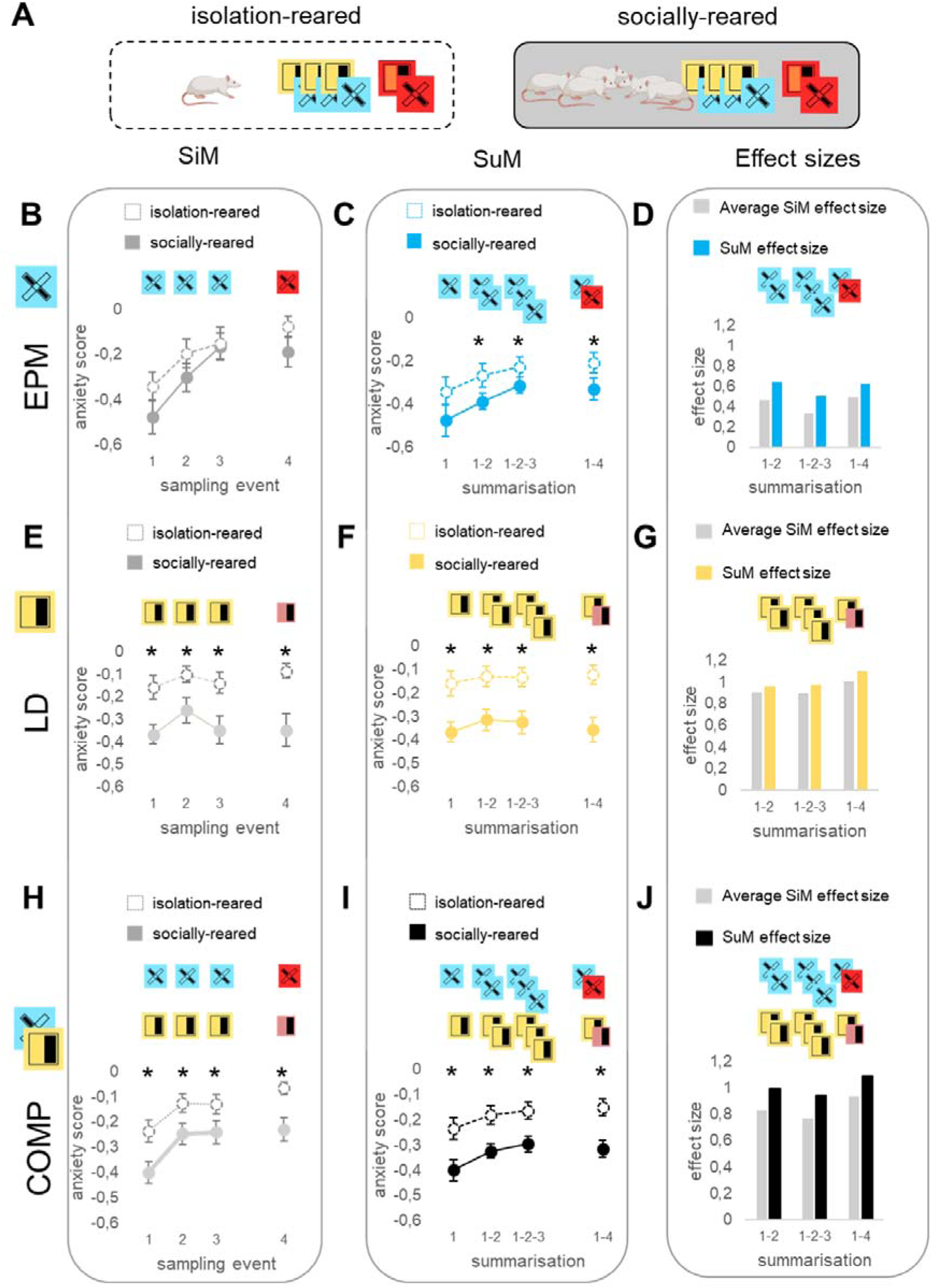
Summary measures as markers of a chronic stress state. **A)** Socially or isolation-reared animals underwent three baseline and one aversive sampling events of the EPM and LD tests then summary measures of different complexity were calculated. **B-E-H)** SiMs (time and frequency-based anxiety scores) were calculated from each sampling events using the EPM (B) or LD tests (E), or from the composite of those (H). **C-F-I)** SiMs or SuMs were calculated from the first or more sampling events (1-2, 1-2-3, 1-4), respectively using the EPM (C), LD (F), or their composites (I). **D-G-J)** The effect sizes of isolation rearing based on averaged sampling events (SuMs) compared to the averages of effect sizes of those very same sampling events without summarization. Briefly, grey and colored bars represent the effect sizes of the comparisons from the first (B-E-H) and the second columns (C-F-I), respectively. All statistical results are presented in Table 1.

### SuMs trace out quantitatively and qualitatively different gene profiles compared to SiMs

Since SuMs represent robust measures of anxiety, we sought to investigate their molecular correlates too. We conducted an RNA-seq experiment on mPFC (exerting the top-down control of anxiety^41,42^) tissue of rats in a resting state. Samples from 27 subjects were chosen based on their anxiety levels. Briefly, we ranked the animals’ COMP scores in ascending order, selecting every second animal for further analysis (Fig.4A). We found that SuMs outperformed SiMs in identifying gene targets, with LD SuMs and EPM SuMs increasing discoveries 3-fold and 8-fold, respectively (Fig.4B). Interestingly, LD-SiM associated genes largely overlap with LD-SuM, while EPM measures correlate with distinct gene sets (Fig.4C). Gene-behavior associations were further characterized by a robustness index showing that most of the molecular targets have a strong relationship with the expression of anxiety measures (Fig.4F).

**Figure 4.**
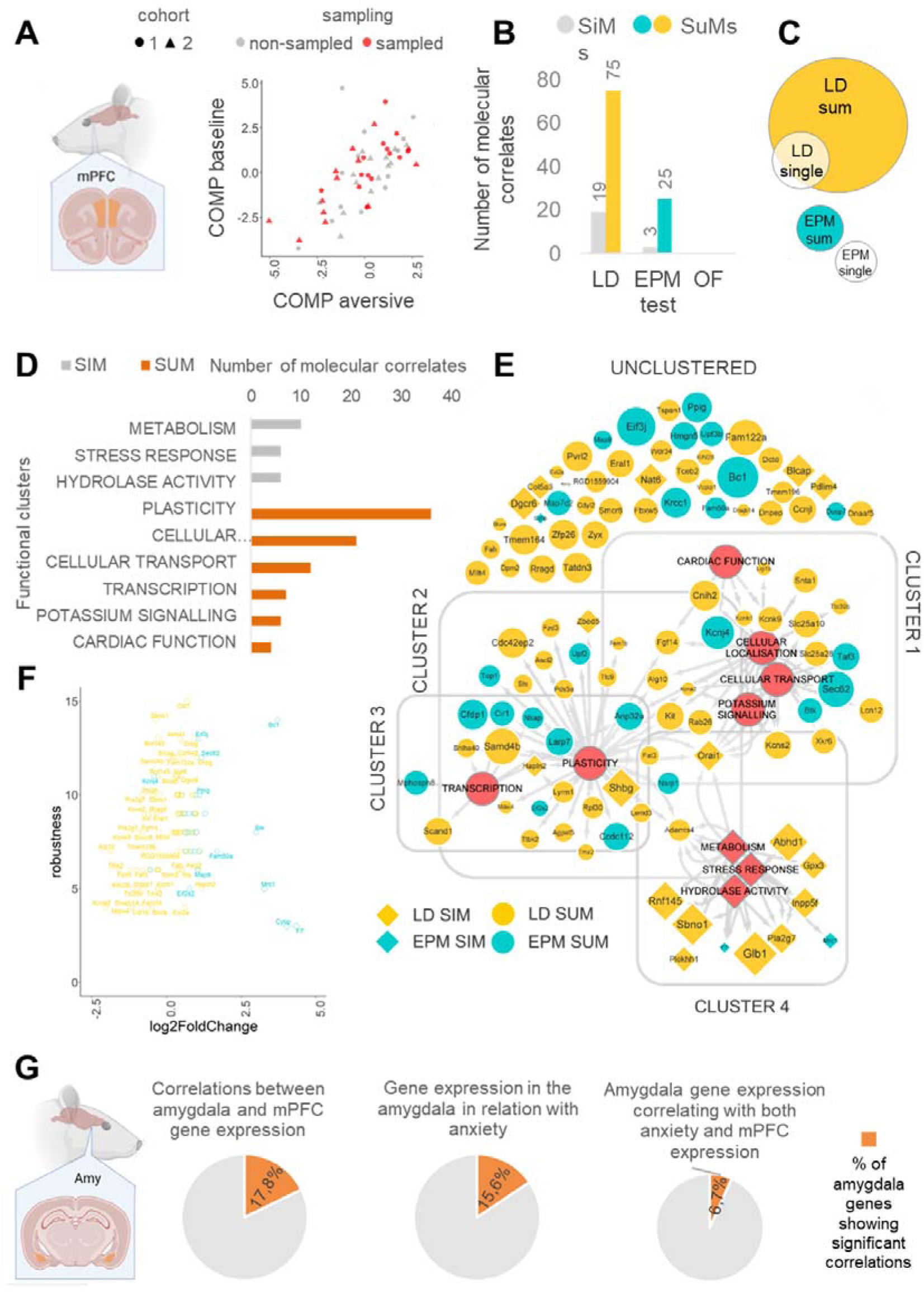
Sensitivity to gene-discovery of differently complex anxiety measures. **A)** left: Representative figure of bilateral mPFC sampling (orange areas). right: Samples for RNA-seq analysis were chosen based on their COMP level **B)** SiM and SuM-associated gene count following FDR correction to multiple comparisons. **C)** Euler diagram representing the proportion of overlaps in gene sets of different sampling approaches. **D) SiM and SuM-associated genes form functionally distinct clusters.** Clusters contain genes with similar annotation profiles based on multiple databases. SiM and SuM-associated genes are grouped into completely non-overlapping clusters defined by metabolism or plasticity, respectively. **E)** Hub-plot of functional annotation of significant genes. Due to the heuristic fuzzy clustering method, a gene does not have to belong to any clusters but can belong to more clusters, while clusters can be determined by multiple functional labels. Cluster 1 includes genes that share many functional labels associated with cardiac function, cellular localization, transport, and potassium signalling. Clusters 2 and 3 are solely characterized by plasticity and transcription functions, respectively. Cluster 4 is characterized by metabolism, stress response, and hydrolase activity-associated functional terms and unites almost exclusively SiM-associated genes. Unclustered genes do not necessarily lack a known function, but do not have the amount of relatedness with enough genes in our pool to form a separate cluster. Gray arrows indicate gene-function associations, while gray lines indicate the borders of clusters. Red circles indicate cluster-determining labels, while color and shape indicate sampling type. The size of hubs is proportional to the correlation coefficient (absolute value) between the gene expression and anxiety. **F)** robustness values of genes as a function of their log2-fold change. **E) Correlations of amygdala gene expressions with PFC gene expression or anxiety variables**. Left piechart: percent of significant correlations between amygdala and mPFC qPCR results. middle: percent of significant correlations between amygdala gene expression and SiMs, SuMs, COMPs or aversive behavior. right: percent of amygdala genes that correlate with both anxiety and mPFC expression. All statistical results are presented in Table 1 and supplementary table 1 and 3.

The distinct gene discovery capabilities of different sampling methods prompted us to investigate their functional variability using the DAVID functional annotation tool for clustering gene discoveries. We found that SiMs correlated with metabolic and stress-related genes, while SuMs established a plasticity-focused gene profile (Fig.4D). The majority of genes associated with SuMs were also putatively involved in cellular localization, cellular transport, potassium signaling, or transcription. Plasticity-associated genes can be further structured to more specific functions, including but not limited to extracellular matrix shaping (ADAMTS4; HAPLN2), general modulation of neuronal development (FGF14; FAT3; FZD3), transcription (BTK; TCEB2; ASCL2; MDM4) or translation (BC1; EIFJ2; EIF2S2; EIF3J; UPF2; UPF3B). Also, a significant percentage of SuM-associated genes are direct effectors of neuronal excitability and neurotransmission, such as KCNK1, KCNK9, KCNA2, KCNS2, and KCNJ4 (Fig.4E). This suggests that our sampling approach not only quantitatively enhances gene discovery, but also yields a qualitatively distinct gene profile.

To confirm our RNASeq findings of differentially expressed genes with a differently sensitive method, we performed a qPCR analysis of the 43 most robust, and the 2 least robust protein-coding genes (Supplementary table 2). When comparing gene expressions between the RNASeq and qPCR data, 17.8% (8) of genes showed significant positive correlations (supplementary table 3). However, 94.6% of the genes that were not significantly correlated between the two different methods showed a fold change value lower than 1.2 (Fig.4F, Supplementary table 1) which is below the typical cut-off criteria for the qPCR analysis^43^. In addition, the remaining 2 non-correlating genes with a fold change higher than 1.2 showed a relatively low expression in the RNASeq and were not detectable with PCR. These were also the 2 least robust genes in our analysis (Supplementary table 2).

Furthermore, we have examined the differential expression of the aforementioned 45 genes in the amygdala, a key region in the regulation of anxiety, and found that after FDR correction, 15.6% (7) of them showed a significant correlation with anxiety (SiMs, SuMs, COMPs or behavior in an aversive environment) (Fig 4G, middle). When examining the correlations between amygdala and mPFC gene expressions measured by qPCR, we found that 17.8% (8) of genes showed a significant positive correlation between the two areas (Fig 4G, left). Furthermore, 6.7% (3) of the investigated amygdala genes showed significant correlations with both anxiety and mPFC gene expression. These genes were Glb1, Pla2g7 and Shbg. These findings suggest that some of the gene targets identified by our method may have a more general role in modulating trait anxiety.

### A consensus score to measure trait anxiety

SuMs provide further predictive value compared to SiMs as a function of their complexity, e.g. the number of sampling points included in those. However, there is an inflection point in every paradigm where the addition of a further sampling point does not significantly enhance predictive value anymore. To offer a standardised approach to measure trait anxiety, we sought to determine a consensus inflection point by comparing the added predictive value of further testing in all paradigms of our study. In Figure 5, we show the improvement multiple testing offers in measured effect sizes of social isolation, correlations between similar anxiety tests under varying aversive conditions, correlations across different anxiety tests, or correlations between anxiety tests and generalized fear responses alongside molecular predictions from our RNASeq analysis. Values of different sampling depths (1, 2 or 3 tests used) are shown as a percentage of the highest prediction value that was achieved in each experiment with a particular test (Fig5B and C) or with any of the tests used (Fig5D). We found that the most easily predictable outcomes were the anxiety variables in aversive conditions, i.e. tests that were done in strong light or following social isolation. Here, a single use of either the EPM or the LD tests could explain above 80% of the maximum prediction value. However, a single EPM could not trace out the effect of isolation, in contrast to the more sensitive EPM SuMs, or an LD SiM that explained approximately the same amount of variability as multiple LD testing. Predicting different contexts was a bigger challenge as it was shown that significant correlations with anxiety in different tests or with freezing in the safe context of the CFP required multiple tests in most of the cases. Finally, finding molecular correlates of anxiety was the most challenging task, as the number of gene correlates continuously increased with every added test occasion, with the most complex SuM variables tracing out the highest number of gene targets in both tests. Finally, comparing the efficacy of the two tests by showing correlations as a percentage of the most accurate predictions of all test variables (Fig.5D) shows that the LD test is superior to the EPM in every paradigm.

**Figure 5.**
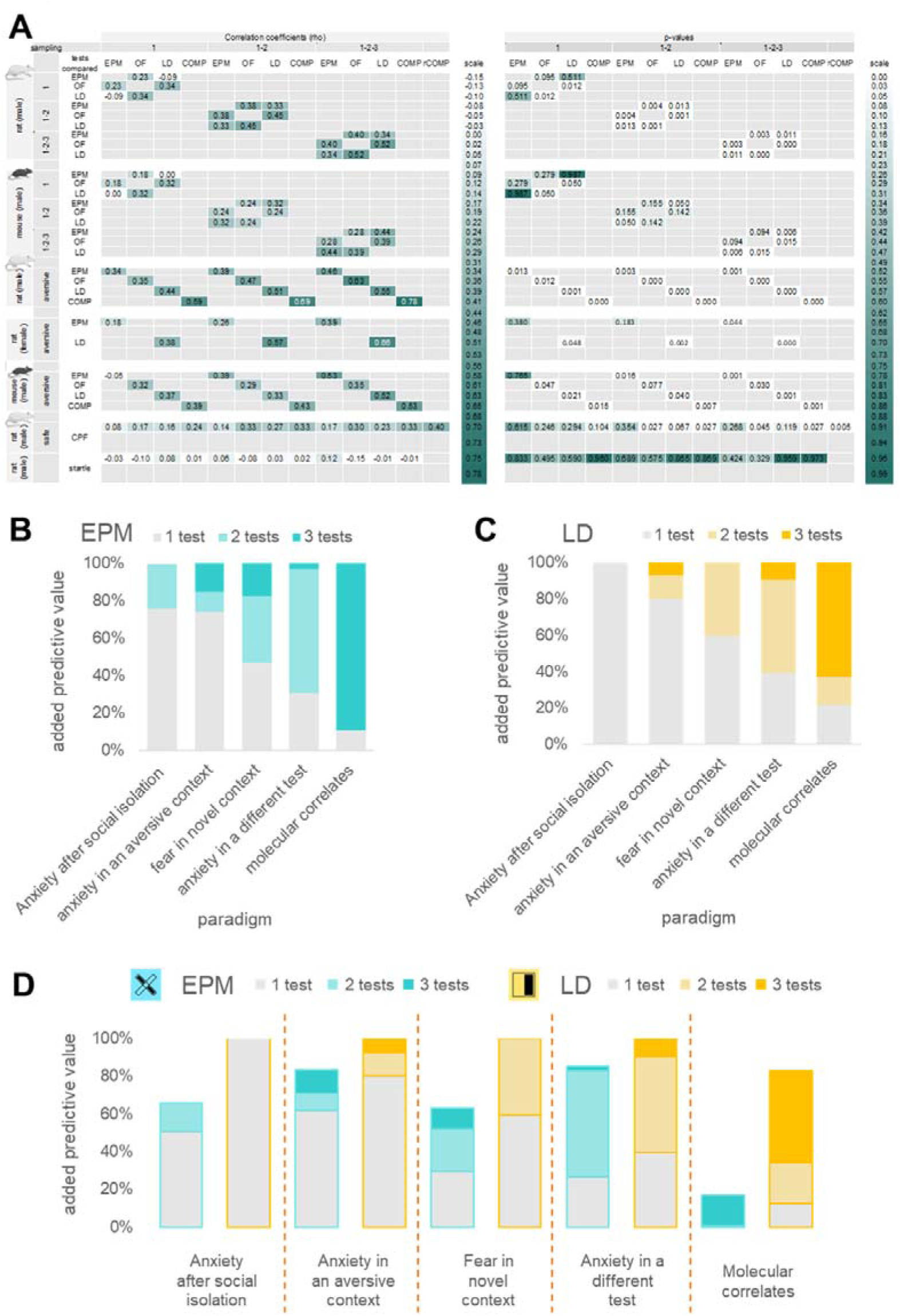
**A) Correlation coefficients and corresponding p-values of the “inter-test correlations” and “behavioral predictions” experiments in rats and mice.** The color code of the cells ranging from white to teal represents the level of correlations or p-values scaled to all parameters and experiments. **B-C) Multiple testing in different paradigms.** Additional predictive value of further sampling points in different paradigms. Values are shown as the percentage of the highest prediction that was reached with a particular test in a particular paradigm. **D)** Additional predictive values are shown as a percentage of the highest prediction that was reached with any of the tests used in a particular paradigm.

## Discussion

Our study aimed to model trait anxiety in rodents to enhance the translational potential of preclinical testing. We hypothesized that the frequently used preclinical anxiety tests measure state anxiety. Since the generally accepted definition of trait anxiety is the tendency of the subject to react with elevated state anxiety in situations^1^, our approach to revealing trait anxiety was to repeat classical anxiety tests multiple times and introduce summary measures of multiple test events and variables.

Here we present the first experimentally validated markers of TA that improve behavioral and molecular predictions of anxiety tests. We found that anxiety tests possess partially overlapping features, which means that creating SuMs is beneficial to explain TA. We validated SuMs in male and female rats and mice as markers of TA by comparing their predictions to classical variables. Since anxiety-like behaviour, habituation, behavioural predictions based on SiM and SuM variables, test differences and test repeatability were similar in males and females, only males were used in the following experiments. We showed that, in proportion with their complexity, SuMs enhance inter-test correlations, strengthen predictions of behavior in threatening environments, and describe the consequences of chronic stress better.

The idea of repeating anxiety tests, especially the EPM has been a topic of debate due to the phenomenon of one-trial tolerance (OTT). OTT refers to two mechanically distinct phenomena; 1) the decreasing level of exploration in the EPM over repeated testing and 2) the decreasing sensitivity of the EPM to measure benzodiazepine (BDZ) effects in repeated testing^44^. The EPM in its original form was developed for assessing BDZ^11^ effects, consequently, any experimental conditions that interfere with this purpose raised concerns about their validity. Therefore, the requirement of BDZ effects has limited the representativity of the EPM test to very few aspects of trait anxiety. Indeed, there is a long list of conditions and drugs that fail to change the behavior expressed in the original EPM protocol, but anxiety-like effects become apparent in different testing conditions, like in more aversive environments, in response to prior stressful experiences, in a modified EPM or through the use of highly anxious animal strains^45–47^. We argue that while the carry-over effect is evident in EPM, additional information can be obtained through multiple testing, as we describe in detail below. Moreover, not all anxiety tests are burdened by a carry-over effect, as behavior in the LD test remains consistent through repetition.

We argue that SuMs represent TA, the shared basis of anxiety-like responses. The TA models of Eysenck, Belzung, and the Spielberger state/trait inventory-based clinical practice align with this concept; all describe SA as an interactional consequence of TA and situational stress^1,3,4,48,49^. These models all agree that the proper description of TA requires multiple sampling in multiple situations; however, our approach is the first that sought to achieve this in preclinical settings. Human phenomena that are explained by TA, such as the clustering of anxiety types in clinical cases or individual-level predictions of future responses are all inconsistent in preclinical models^4,16,18^. In our study, the gradual improvement in inter-test correlations and predictive power when increasing the complexity of sampling supports that SuMs converge to TA. To our best knowledge, this is the first TA paradigm that modifies experimental analysis instead of experimental subjects^20–22,50^, giving way to individual phenotyping on a level that the time-and effort-consuming experiments of modern neuroscience desire.

Next, we aimed to physiologically validate SuMs as measures of TA, so we compared these with SiMs in describing whole-genome expression profiles in mPFC samples. The use of SuMs revealed a quantitatively and qualitatively different genetic profile from SiMs and enhanced gene discoveries up to an 8-fold improvement. Higher discovery rates of SuMs indicate they are more sensitive to detect gene-anxiety associations. The reason for that might be that SuMs describe a greater proportion of variance in gene expression data of the same subject. This allows us to use analytical approaches which require higher statistical power but also possess greater biological relevance, such as describing gene expression with continuous anxiety scores instead of categorical anxiety variables like high and low anxiety groups. Also, standard designs require fewer subjects to get valuable data which aligns with the 3R principle. Finally, greater sensitivity improves the discovery rate of new gene targets of anxiety^51^.

Furthermore, genes detected with SuMs clustered into functionally different groups than SiMs indicating that besides having superior sensitivity, SuMs explain a previously undescribed type of variability. These previously undiscovered gene products are all potential candidates for the treatment of trait anxiety and thus anxiety disorders. We have also tested the robustness of the molecular findings, to further increase the translatability of our results.

SuM genes show high functional relatedness and are associated with neural plasticity and psychiatric disorders through different mechanisms and pathways. SuMs are positively correlated with the expression-levels of perineuronal net (PNN) and perinodal extracellular matrix (ECM) shaping molecules that influence membrane currents, excitability, and synaptic reorganization and were described in mental disorders^52^. ADAMTS4 is responsible for the cleavage of PNN elements^52,53^, while HAPLN2 is an ECM composer that establishes the proper conductivity of neurons by stabilizing Ranvier nodes^52^. SuMs are negatively correlated with the trophic factor FGF14 that establishes proper excitability of neurons^54,55^. SuMs correlate positively with many general modulators of gene transcription and translation that control the expression of wide clusters of genes. Examples of this are Bruton tyrosine kinase (BTK), the up-frameshift factors UPF2 and UPF3b^56^, Elongin-B (TCEB2)^57^, brain cytoplasmic RNA 1 (BC1)^58^, or subunits of the Eukaryotic translation initiation factors EIF2s2, EIFJ2, and EIF3J^59,60^. Finally, we found multiple types of differently regulated potassium channels, which are direct modulators of membrane currents. This indicates that trait anxiety correlates with a consistently altered gene expression, excitability, and the capability of neuronal reorganization in resting conditions in the mPFC. This data represents a range of promising candidates that could trace out novel interventions to treat anxiety.

To propose a versatile tool for measuring TA, we compared the predictive value of SiMs and SuMs of different tests in all the paradigms used in this study. We found that predicting behavior or molecular correlates varies in difficulty across paradigms. While we can get accurate predictions using a single test when predicting responses to an aversive environment, we need at least two tests for predicting behavior in a different context, and all three tests when searching for molecular correlates of anxiety. None of the variables, regardless of their complexity were able to predict acoustic startle, which shows that SuMs predicting future anxious states stronger than SiMs is not due to technical factors, but rather due to SuMs predicting specific emotional domains. Our analysis showed that the LD test outperformed the EPM in every paradigm, with a single LD test providing as much insight as multiple EPM tests, while two LD tests achieved 96% of the most accurate predictions. Additionally, LD SuMs identified four-times more anxiety-related genes than any EPM variable. Based on these findings, we recommend conducting at least two LD tests with an inter-test interval of one day, as an effective and precise approach for measuring rodent trait anxiety.

In summary, here we provide a novel sampling and analysis pipeline that enables us to measure TA in preclinical research. Our paradigm is based on an extended sampling process using popular anxiety tests, consequently making it applicable to any model and feasible to any laboratory. We argue that the correlation between the complexity of sampling and the amount of information gained of the anxious phenotype of animals is robust and representative to various conditions. Such correlation was confirmed by applying multiple testing conditions: in baseline and aversive testing environment, in multiple test types, in naïve as well as chronic stress-exposed animals, and through using different cohorts, species and sexes of subjects. In line with this, the molecular profiling of TA in female rodents is a crucial further step to make due to potential differences at the level of the mPFC^61^. Using SuMs in experiments boosts behavioral and molecular phenotyping and predictions, consequently reducing the minimally sufficient sample sizes while maximizing the discovery rate of novel treatment candidates. In addition, SiMs reveal a distinct, plasticity-focused gene profile associated with trait anxiety. We encourage colleagues to adopt this detailed phenotyping approach, potentially bridging the gap between preclinical and clinical anxiety research.

## Materials and Methods

Detailed methods are provided in the supplementary materials.

### Animals

Six cohorts of animals were used throughout this study. After arrival at the facility, all cohorts were left to adapt to the housing conditions for 2 weeks. Two cohorts of adult male Wistar rats (Charles-River, n=54, age= 100 days) were used to examine trait anxiety and its neural correlates. To assess replicability, behavioral assessment was repeated with adult female Wistar rats (Charles-River, n=27, age= 130 days), and adult male C57BL/6J mice (Jackson Laboratory, n=40, age= 95 days). One cohort of adult male Long-Evans rats (Charles-River, n=27, age= 110 days) was used to explore the correlations between acoustic startle responses, generalized fear behavior, and trait anxiety in the conditioned fear paradigm (CFP). Additionally, one cohort of adult male Wistar rats (Charles-River, n=30, age at testing= 180 days) was used to investigate the effects of social isolation on trait anxiety. Sample sizes for the first rat experiment (n=54) were calculated using the pwr R package aiming to measure a minimum r=0.35 spearman linear correlation coefficient at alpha=0.05 significance level and beta=0.8 power. Following the first experiment we decreased sample sizes according to the robustness indices of behavior-behavior and behavior-gene expression correlations.

### Experimental design

Wistar rats were tested according to a semi-randomized testing protocol for 3+2 weeks (Fig.1). The test battery involved a 3-week behavioral sampling with the most commonly used anxiety tests; the Elevated Plus-Maze^10,11^ (EPM), Open Field^14,24^ (OF), and Light-Dark test^12,13^ (LD), each repeated three times for all individuals. Test order and time-of-day biases were eliminated by creating six different test combinations through semi-randomization of test orders, with an equal number of animals assigned to each combination. After the initial 3-week testing period, animals underwent an additional week of testing in a more aversive experimental condition within a brightly lit test arena. Two days after the aversive week, a different type of open field test (OF2) was performed as the final test for all animals. For the social isolation protocol, upon weaning at P21, animals were reared in groups of 4 (social), or alone in a cage (isolated) until testing in adulthood. Female rats, male rats in the social isolation cohort and mice underwent the 3-week baseline anxiety testing, and the additional week of testing in aversive conditions. Long-Evans rats first underwent a 3-week anxiety testing protocol, followed by an acoustic startle paradigm 3 days later, and finally a fear conditioning paradigm (CFP) a week after that. For the experiments described above, animals were randomly chosen for each group. For anxiety tests, time spent in the aversive zone, frequency, and latency of aversive zone entries were analyzed automatically using Noldus EthoVision XT1541, head-dips (HD) and rearing (R) were analyzed manually using Solomon Coder^24^ by experimenters that were blind to experimental groups.

### Behavioral measures

To attenuate the possibly state anxiety-driven fluctuation of behavioral variables across repeated testing, therefore to potentially detect stable traits, we introduced summary measures of behavior. Our enhancements to the conventional preclinical approach involved two key aspects: 1) Using the outcomes of the principal component analysis shown in Figure 2C, we calculated z-scores by summing variables that share the same dimension. Before summation, we scaled these variables by subtracting the population mean from each individual’s values, then dividing by the standard deviation. SiMs and SuMs were measured in the same animals. We refer to the scaled behavioral variables characterized in the first tests (EPM, OF, LD performed during the first week of testing) as single measures (SiMs). 2) By averaging the scaled variables of the 2-, or 3-time repetition of each anxiety test type, we created summary measures (SuMs). To assess the effectiveness of SuMs in capturing the common underlying factors across different anxiety tests, we compared the strength of individual-level correlations using SiMs versus SuMs. Stronger inter-test correlations indicate that a variable better captures the shared background, potentially reflecting trait anxiety. Composite z-scores (COMP) were created by averaging the SiMs or SuMs of different tests. Means and standard errors are reported for behavioural measures.

### Statistical analysis

was done in R statistical environment^25^. Principal component analysis (‘factoextra’ package^26^) of all analyzed behavioral variables from the first week of testing was performed to investigate the shared variance between different behavioral variables. For correlations, Spearman correlation analysis (‘Hmisc’ package^27^) was used. The analysis of test-repeatability (‘rptr’ package^28^) was done for each test-type as described previously^29^, by calculating the proportion of within-test variance divided by total variance^30^. False discovery rate (FDR) correction for multiple comparison testing was done using the Benjamini-Hochberg method^31^.

### RNASeq analysis

Microdissection of bilateral medial prefrontal cortex (mPFC) tissue was performed 2 weeks after the last experiment (OF2). Whole transcriptome-analysis via RNASeq was performed on the homogenized mPFC tissue. RNA sequencing was performed on 27 animals, chosen by their COMP scores (Fig.5A). Isolation and mRNA library preparation was carried out by the Department of Biochemistry and Molecular Biology at University of Debrecen, using the NEBNext Ultra II RNA Library Prep with Purification Beads Kit (New England Biolabs). Single-end RNA sequencing was performed on an Illumina NextSeq500 sequencer. Regression analysis to reveal differentially expressed genes (DEGs) was done using the R Bioconductor package DESeq2^32^. Analysis of gene-behavior regressions was carried out using a negative binomial generalized linear model, with anxiety-scales generated from behavior in different test-types (EPM/OF/LD) across test repetitions (SiM/SuM) as continuous covariates in the design. False discovery rate (FDR) correction was done using the Independent Hypothesis Weighting method^31^. Only genes with FDR corrected p-values below 0.05 were considered to show significant differential expression. Functional and pathway analysis of detected significant DEGs was done through the online functional annotation tool DAVID^33,34^. Connections were visualized using Cytoscape^35^. Cluster-determining functions, merging and associated genes are shown in Supplementary table 2.

### qPCR analysis

We performed qPCR analysis of total RNA from mPFC and AMY. For mPFC, the RNA sample isolated for RNASeq was used for further analysis. Bilateral microdissection of the amygdala was performed as described for mPFC tissue. Gene expression patterns were analyzed using custom 384-well TaqMan Gene Expression Array Cards (Applied Biosystems, USA). These cards measured the expression of 45 genes of interest, which were selected based on the robustness of mPFC RNA sequencing experiments. Inter-run calibration values were calculated based on the run control samples, using Actb and Gapdh as reference genes, as per the method described by Hellemans^36^. The normalized relative gene expression values were calculated using the 2-ΔΔCT method^37^. These values were further normalized to the inter-run calibration values. Expression values with threshold cycles greater than 35 were excluded from the analyses.

### Robustness

We characterised the stability of behaviour-behaviour and behaviour-gene expression correlations with the robustness index. The robustness index gives the smallest number of randomly excludable samples that likely impair the significance of such a relationship. The index is calculated by a custom-written script that reanalyses the Spearman correlations in randomly selected, different sample size sub-cohorts of the original population. We made 20000 correlations on all possible sample size sub-cohorts and calculated minimums, maximums, standard deviations and means of their p-values. We determined the cutoff criteria of a robustness index as the smallest sample size in which the standard deviation of the correlations’ p-values reaches the 0.05 alpha level. The function in R is available upon request.

## Data Availability Statement

Raw count values of RNA sequencing data and the corresponding behavioural data is publicly available at the Zenodo online data repository through the accession number and doi 10.5281/zenodo.14236344.

## Acknowledgement

This work was supported by the Hungarian Scientific Research Fund (Grant No. K125390), Eötvös Loránd Research Network (Grant No. SA-49/2021), Hungarian Brain Research Program (Grant No. 2017-1.2.1-NKP-2017-00002), for dr. Éva Mikics, and the Bolyai Janos Research Fellowship for Mate Toth. We would like to thank Dr. Kornél Demeter, the head of the Unit of Behavioral Studies at IEM for providing the environment for keeping laboratory animals and conducting experiments. On behalf of the "NGS analysis in neurological and behavioral sciences" project we are grateful for the possibility to use ELKH Cloud (see Héder et al. 2022; https://science-cloud.hu/) which helped us achieve the results published in this paper.

