## supplementary FIGURES for "Improving anxiety research: novel approach to reveal trait anxiety through summary measures of multiple states"

#### Supplementary Figure1

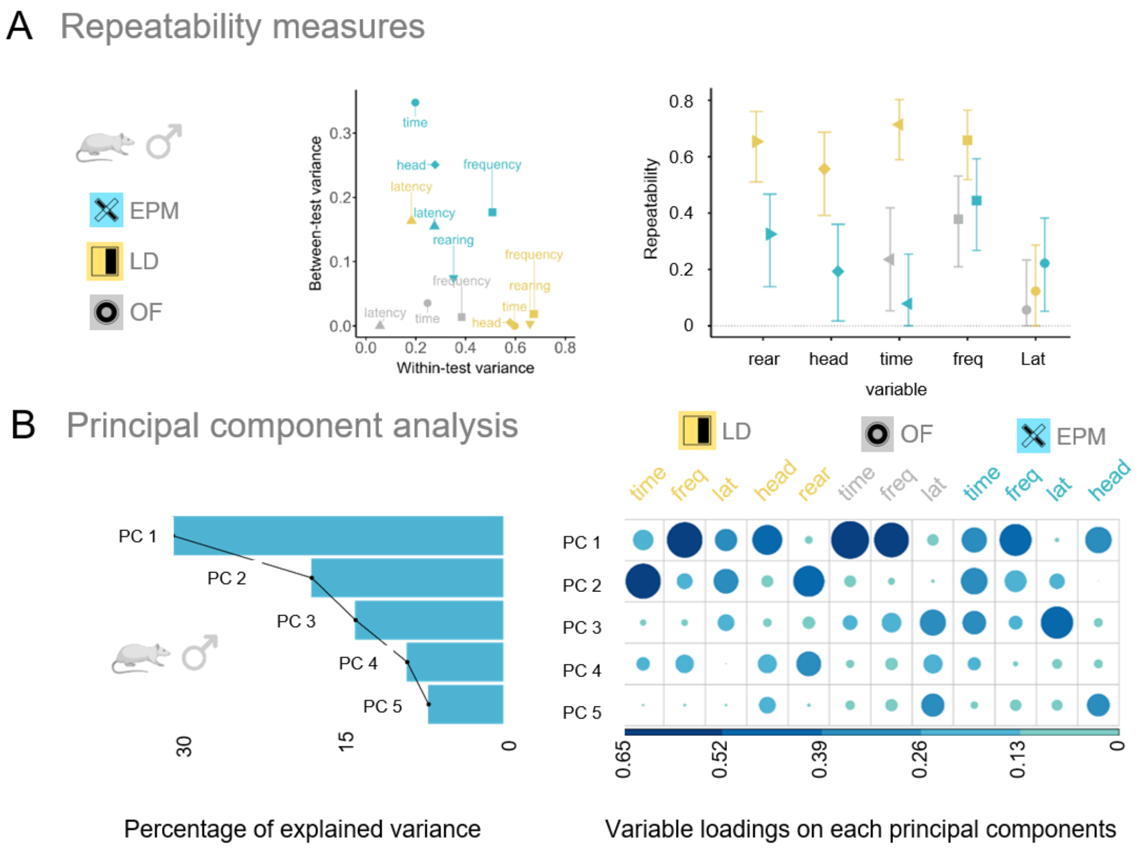


**Supplementary Figure 1.** Shared and unique features of anxiety tests in male rats. **A) left:** all measured variables in variable space defined by a between-test / within-test variance. **right:** bootstrap analysis-based repeatability scores of variables with 95% confidence intervals (represented by error bars). Variables with confidence intervals not including a 0 repeatability score are considered repeatable. **B) left:** percentage of explained variance by the first 5 principal components (PC) of a PCA analysis on a single (first) anxiety test. **right:** variable representations on each PC. Size and color hue is proportional to the level of representation of a given loading. **Abbreviations:** EPM: elevated plus-maze test, OF: open field test, LD: light/dark test, Lat: latency to the aversive compartment (AC)(sec), freq: enter frequencies to AC (count), time: %time spent in AC, head: head dipping from the platform, rear: rearing behavior.

### Supplementary Figure 2

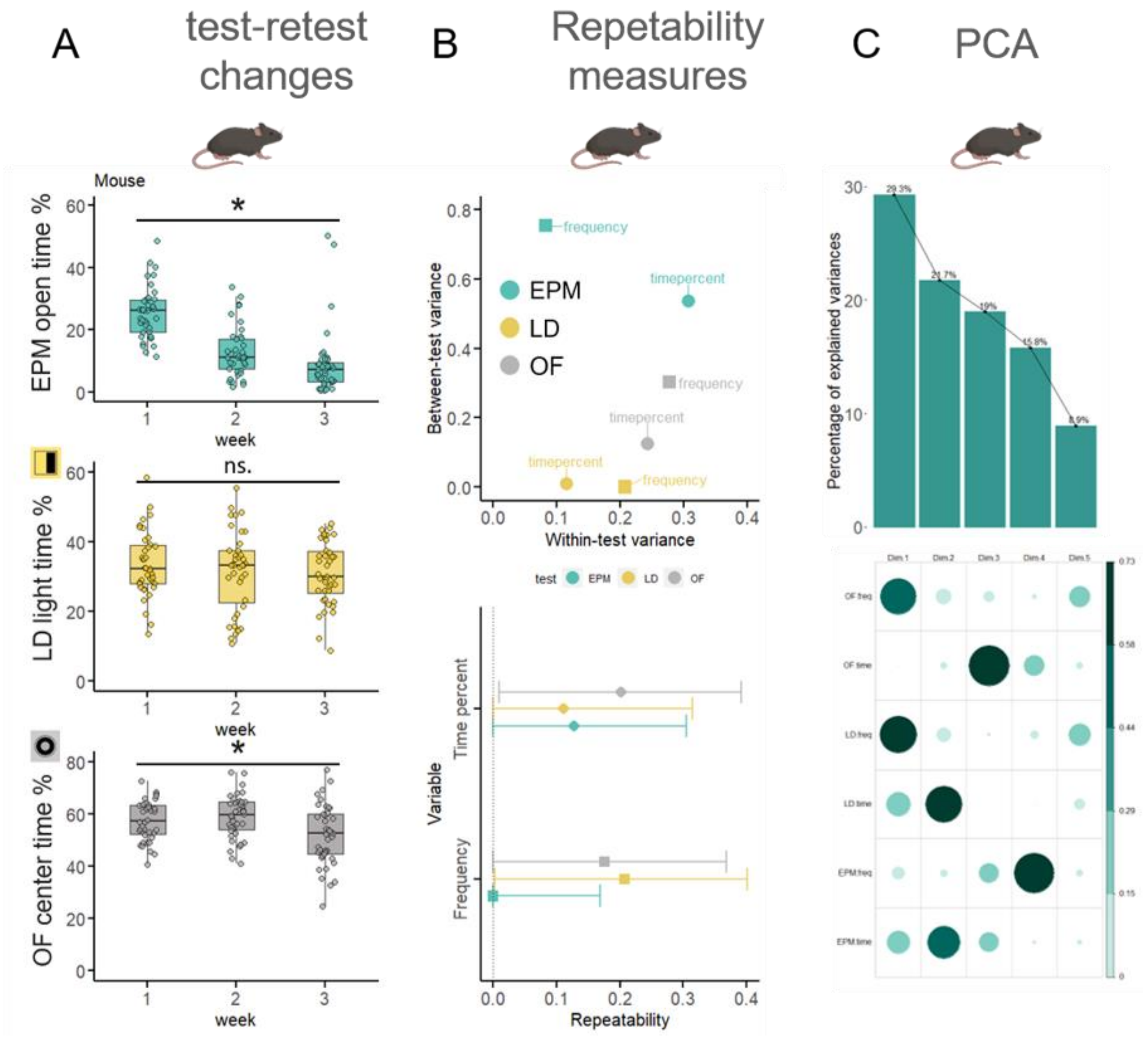


**Supplementary Figure 2.** Shared and unique features of anxiety tests in mice. **A)** The effect of test repetitions on the most frequently used anxiety-like behavioural outcome (% of time spent in the aversive compartment) of the EPM, LD and OF anxiety tests from top to bottom, respectively. **B)** top: all measured variables in variable space defined by a between-test / within-test variance. bottom: bootstrap analysis-based repeatability scores of variables with 95% confidence intervals (represented by error bars). Variables with confidence intervals not including 0 repeatability score are considered repeatable. **C)** top: percentage of explained variance by the first 5 principal components (PC) of a PCA analysis on a single (first) anxiety test. bottom: variable representations on each PC. Size and colour hue is proportional to the level of representation of a given loading. Abbreviations: EPM: elevated plus-maze test, OF: open field test, LD: light/dark test, Lat: latency to the aversive compartment (AC)(sec), freq: enter frequencies to AC (count), time: %time spent in AC, head: head dipping from the platform, rear: rearing behaviour.

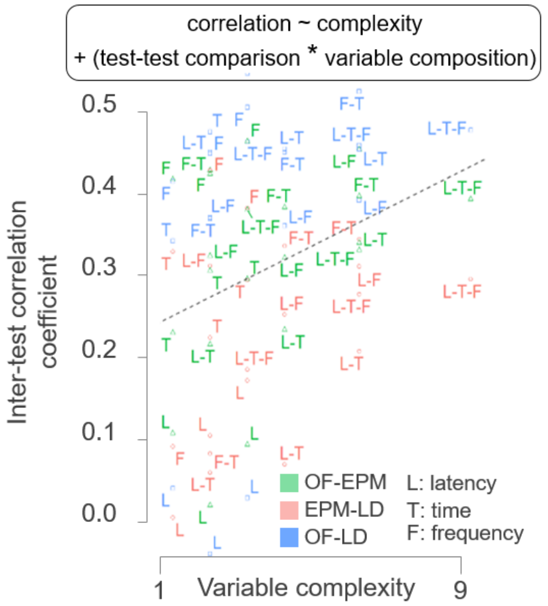


**Supplementary figure 3. Inter-test correlations as a function of variable complexity. A)** Inter-test correlations between increasingly complex anxiety variables – SiMs and SuMs – were plotted in response to the number of test types, events, and variables that were included in each measure. According to our final statistical model, inter-test correlations were significantly influenced by variable complexity as a fixed effect, and variable composition and test composition as interacting random effects. The color code indicates different test-test comparisons, while the abbreviations indicate which variables were used to compose a SuM. L: latency, T: time, F: frequency.

### Supplementary Figure 4

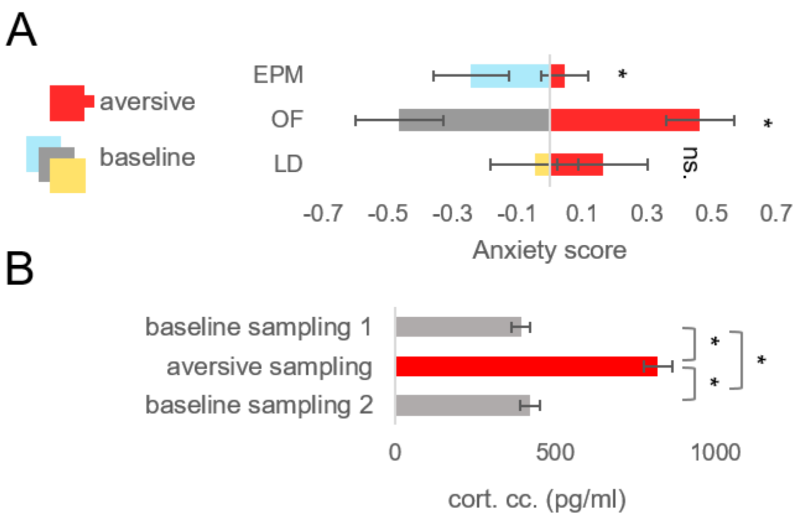

**Supplementary figure 4. A)** Anxiety score from the last baseline and the aversive sampling**. B)** Blood plasma corticosterone levels in resting state before (1) and after (2) the test battery, and immediately following aversive anxiety sampling.

#### Supplementary Figure 5

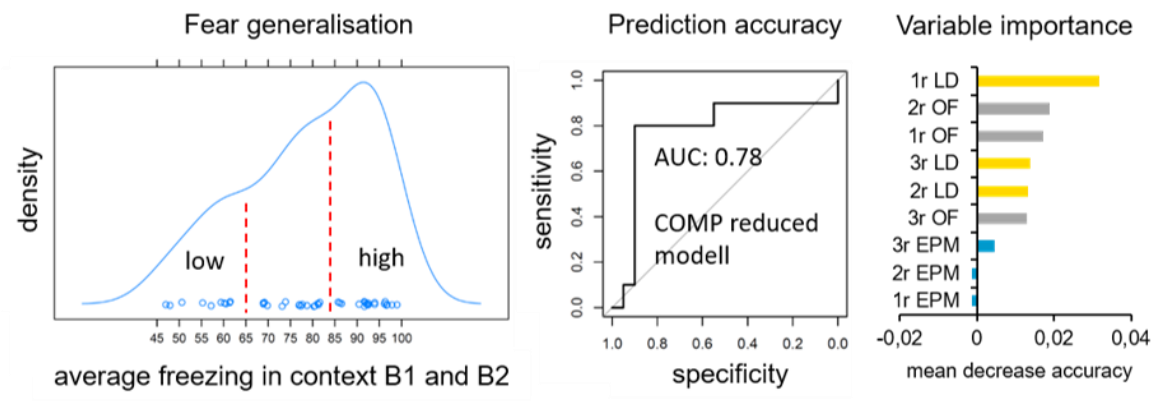


**Supplementary Figure 5.** Random Forest classification of low and high freezer subgroups. **Left**: The freezing behaviour of rats in a neutral, safe context (context B1 and B2) were measured following fear conditioning. Animals were allocated to low and high freezer groups based on the density of averaged freezing. **Middle**: Prediction accuracy was calculated from the ratio of true positive (sensitivity) and true negative (specificity) votes of our reduced COMP model consisting of 1 LD and 2 OF variables. **Right**: Reduced models were developed based on variable importance scores of a saturated model.

| model | sampling | gene symbol | base means | $log_2$ FC | $log_2$ FC SE | robustness | FDR-adjusted p-value | IHW weight |
|---|---|---|---|---|---|---|---|---|
| EPM | SiM | Cytip | 23.606 | 4.028 | 0.960 | 3 | 0.050 | 2.771 |
| | | F7 | 22.638 | 4.342 | 1.022 | 3 | 0.050 | 2.771 |
| | | Mrc1 | 155.711 | 3.277 | 0.780 | 5 | 0.050 | 4.962 |
| | SuM | Anp32a | 1803.473 | 0.481 | 0.113 | 9 | 0.013 | 2.500 |
| | | Bc1 | 25643.881 | 3.720 | 0.568 | 14 | 0.000 | 2.447 |

| | | | | | | | |
|---|---|---|---|---|---|---|---|
| | Btk | 38.921 | 2.995 | 0.763 | 8 | 0.045 | 1.325 |
| | Ccdc112 | 176.938 | 0.929 | 0.218 | 9 | 0.026 | 0.650 |
| | Cfdp1 | 806.939 | 0.499 | 0.126 | 10 | 0.026 | 2.500 |
| | Cir1 | 476.013 | 0.870 | 0.191 | 8 | 0.004 | 2.500 |
| | Dusp7 | 851.688 | -0.534 | 0.130 | 6 | 0.016 | 3.026 |
| | Eif2s2 | 1070.662 | 0.767 | 0.205 | 5 | 0.045 | 2.500 |
| | Eif3j | 696.432 | 1.062 | 0.205 | 13 | 0.001 | 1.200 |
| | Fam50a | 167.672 | 1.704 | 0.434 | 7 | 0.045 | 1.200 |
| | Hmgn5 | 110.513 | 1.294 | 0.299 | 9 | 0.022 | 0.673 |
| | Kcnj4 | 908.471 | -0.407 | 0.110 | 11 | 0.045 | 3.026 |
| | Krcc1 | 239.564 | 0.999 | 0.230 | 10 | 0.021 | 0.673 |
| | Larp7 | 525.165 | 0.753 | 0.162 | 9 | 0.003 | 2.500 |
| | Map7d2 | 1219.487 | 0.674 | 0.137 | 8 | 0.001 | 2.500 |
| | Map9 | 582.845 | 1.049 | 0.270 | 6 | 0.033 | 2.500 |
| | Mphosph8 | 1155.360 | 0.727 | 0.157 | 8 | 0.003 | 2.500 |
| | Nkap | 140.282 | 1.109 | 0.257 | 7 | 0.016 | 1.200 |
| | Nsrp1 | 294.536 | 0.949 | 0.217 | 7 | 0.020 | 0.627 |
| | Ppig | 1241.829 | 1.107 | 0.196 | 10 | 0.000 | 2.500 |
| | Sec62 | 1834.781 | 1.166 | 0.207 | 12 | 0.000 | 2.447 |
| | Taf3 | 336.897 | 0.595 | 0.143 | 9 | 0.035 | 0.673 |

| | | | | | | | | |
|---|---|---|---|---|---|---|---|---|
| | | Top1 | 420.345 | 0.825 | 0.192 | 7 | 0.017 | 1.200 |
| | | Upf2 | 529.868 | 0.623 | 0.142 | 7 | 0.008 | 2.500 |
| | | Upf3b | 574.895 | 0.998 | 0.246 | 8 | 0.046 | 0.627 |
| LD | SiM | Abhd1 | 145.172 | 0.460 | 0.112 | 13 | 0.034 | 0.999 |
| | | Adamts4 | 78.607 | 0.652 | 0.165 | 9 | 0.049 | 1.288 |
| | | Blcap | 1577.023 | 0.291 | 0.065 | 11 | 0.024 | 0.449 |
| | | Col5a3 | 326.879 | 0.586 | 0.149 | 8 | 0.026 | 4.378 |
| | | Dgcr6 | 933.387 | 0.385 | 0.093 | 11 | 0.033 | 1.062 |
| | | Glb1 | 160.858 | 0.857 | 0.153 | 15 | 0.000 | 1.288 |
| | | Gpx3 | 181.445 | 0.983 | 0.234 | 9 | 0.024 | 1.464 |
| | | Hapln2 | 334.366 | 1.182 | 0.288 | 7 | 0.031 | 1.288 |
| | | Inpp5f | 699.704 | 0.417 | 0.095 | 10 | 0.019 | 1.046 |
| | | Nat6 | 285.529 | 0.393 | 0.092 | 11 | 0.024 | 1.288 |
| | | Orai1 | 104.910 | 0.963 | 0.225 | 10 | 0.026 | 0.789 |
| | | Pdlim4 | 397.688 | 0.410 | 0.107 | 9 | 0.031 | 4.313 |
| | | Pla2g7 | 184.910 | -0.932 | 0.209 | 10 | 0.018 | 0.999 |
| | | Plekhh1 | 406.534 | 0.988 | 0.235 | 8 | 0.015 | 4.378 |
| | | Rmrp | 40.122 | -6.810 | 1.370 | 0 | 0.004 | 1.288 |
| | | Rnf145 | 1136.341 | -0.283 | 0.059 | 13 | 0.004 | 1.244 |
| | | Sbno1 | 1076.216 | -0.406 | 0.081 | 14 | 0.004 | 0.825 |

| | | | | | | | | |
|---|---|---|---|---|---|---|---|---|
| | | Shbg | 53.145 | 1.004 | 0.207 | 13 | 0.004 | 1.252 |
| | | Zbed5 | 428.279 | 0.362 | 0.099 | 8 | 0.048 | 4.313 |
| | SuM | Adamts4 | 78.607 | 0.570 | 0.159 | 7 | 0.047 | 1.587 |
| | | Agpat5 | 483.012 | -0.263 | 0.070 | 6 | 0.035 | 1.652 |
| | | Alg10 | 32.986 | -1.880 | 0.460 | 7 | 0.015 | 1.652 |
| | | Ascl2 | 66.192 | 1.032 | 0.245 | 6 | 0.013 | 1.959 |
| | | Bhlhe40 | 1299.835 | -0.301 | 0.080 | 6 | 0.029 | 2.396 |
| | | Blcap | 1577.023 | 0.291 | 0.057 | 12 | 0.004 | 0.640 |
| | | Blvra | 296.247 | -0.473 | 0.133 | 4 | 0.044 | 2.044 |
| | | Ccnjl | 34.154 | 0.969 | 0.225 | 9 | 0.011 | 1.652 |
| | | Cdc42ep2 | 202.890 | 0.374 | 0.095 | 10 | 0.023 | 1.652 |
| | | Cdyl2 | 59.969 | -0.790 | 0.210 | 6 | 0.034 | 1.959 |
| | | Cnih2 | 2437.705 | 0.352 | 0.076 | 10 | 0.004 | 2.396 |
| | | Col5a3 | 326.879 | 0.634 | 0.127 | 12 | 0.004 | 1.448 |
| | | Dctd | 48.110 | 0.596 | 0.166 | 8 | 0.047 | 1.652 |
| | | Dgcr6 | 933.387 | 0.361 | 0.087 | 10 | 0.021 | 0.847 |
| | | Dnaaf5 | 117.892 | 0.461 | 0.128 | 8 | 0.046 | 1.476 |
| | | Dnajb14 | 67.919 | -1.098 | 0.293 | 4 | 0.042 | 1.120 |
| | | Dnpep | 964.720 | 0.254 | 0.067 | 8 | 0.044 | 0.913 |
| | | Dpm2 | 315.731 | 0.369 | 0.102 | 6 | 0.045 | 1.589 |

| | | | | | | |
|---|---|---|---|---|---|---|
| Eral1 | 351.824 | -0.308 | 0.084 | 9 | 0.037 | 2.044 |
| Evi2a | 277.589 | 0.618 | 0.168 | 4 | 0.042 | 1.448 |
| Fah | 200.247 | 0.497 | 0.122 | 6 | 0.015 | 1.652 |
| Fam122a | 214.924 | 0.348 | 0.088 | 12 | 0.024 | 1.549 |
| Fat3 | 152.133 | -1.018 | 0.277 | 6 | 0.042 | 1.476 |
| Fbxw5 | 737.113 | 0.287 | 0.070 | 8 | 0.037 | 0.347 |
| Fem1b | 221.242 | -0.608 | 0.172 | 4 | 0.049 | 1.698 |
| Fgf14 | 31.569 | -0.802 | 0.221 | 8 | 0.042 | 1.834 |
| Fzd3 | 55.025 | -1.216 | 0.277 | 6 | 0.009 | 1.959 |
| Glb1 | 160.858 | 0.699 | 0.160 | 10 | 0.010 | 1.698 |
| Gpx3 | 181.445 | 0.974 | 0.213 | 9 | 0.006 | 1.652 |
| Hapln2 | 334.366 | 1.080 | 0.272 | 5 | 0.023 | 1.448 |
| Kcna2 | 36.710 | -2.040 | 0.545 | 4 | 0.035 | 1.959 |
| Kcnk1 | 1531.718 | -0.273 | 0.077 | 5 | 0.042 | 2.396 |
| Kcnk9 | 52.273 | -1.220 | 0.334 | 8 | 0.042 | 1.834 |
| Kcns2 | 60.124 | -0.745 | 0.180 | 9 | 0.014 | 1.959 |
| Kit | 183.256 | -0.620 | 0.148 | 9 | 0.014 | 1.652 |
| Klhl28 | 38.999 | -1.140 | 0.323 | 5 | 0.048 | 1.834 |
| Lcn12 | 37.886 | 0.785 | 0.208 | 8 | 0.035 | 1.652 |
| Lemd3 | 277.339 | -0.322 | 0.088 | 6 | 0.042 | 1.589 |

| | | | | | | | |
|---|---|---|---|---|---|---|---|
| | Lrp1b | 352.926 | -0.844 | 0.236 | 4 | 0.049 | 1.409 |
| | Lyrm1 | 182.250 | 0.463 | 0.126 | 7 | 0.042 | 1.698 |
| | Mdm4 | 95.155 | -1.173 | 0.319 | 4 | 0.042 | 1.587 |
| | Mllt4 | 1188.392 | -0.247 | 0.065 | 8 | 0.042 | 0.868 |
| | Nat6 | 285.529 | 0.341 | 0.089 | 8 | 0.027 | 2.044 |
| | Orai1 | 104.910 | 0.908 | 0.212 | 9 | 0.013 | 1.476 |
| | Pds5a | 192.446 | -0.572 | 0.156 | 6 | 0.042 | 1.549 |
| | Pla2g7 | 184.910 | -0.807 | 0.205 | 8 | 0.024 | 1.549 |
| | Pvrl2 | 86.300 | 0.495 | 0.132 | 10 | 0.037 | 1.597 |
| | Rab26 | 1010.801 | 0.305 | 0.081 | 8 | 0.044 | 0.847 |
| | RGD1559904 | 298.985 | -0.327 | 0.091 | 7 | 0.045 | 1.652 |
| | Rnf145 | 1136.341 | -0.268 | 0.054 | 11 | 0.004 | 0.347 |
| | Rpl30 | 472.323 | 0.662 | 0.164 | 7 | 0.019 | 1.507 |
| | Rragd | 188.492 | -0.452 | 0.107 | 9 | 0.013 | 1.698 |
| | Samd4b | 829.421 | 0.215 | 0.051 | 12 | 0.029 | 0.347 |
| | Sbno1 | 1076.216 | -0.347 | 0.081 | 10 | 0.014 | 0.847 |
| | Scand1 | 644.680 | 0.484 | 0.115 | 9 | 0.014 | 1.589 |
| | Shbg | 53.145 | 0.942 | 0.195 | 12 | 0.004 | 1.652 |
| | Slc25a10 | 322.184 | 0.392 | 0.095 | 9 | 0.015 | 1.448 |
| | Slc25a28 | 536.981 | 0.295 | 0.071 | 8 | 0.015 | 1.507 |

| | | | | | | | |
|---|---|---|---|---|---|---|---|
| | Smcr8 | 96.412 | -0.931 | 0.227 | 8 | 0.015 | 1.658 |
| | Snta1 | 641.738 | 0.223 | 0.062 | 8 | 0.047 | 1.467 |
| | Sts | 173.874 | 0.531 | 0.147 | 6 | 0.044 | 1.698 |
| | Tatdn3 | 213.583 | 0.528 | 0.113 | 10 | 0.004 | 1.698 |
| | Tceb2 | 829.548 | 0.475 | 0.112 | 8 | 0.015 | 0.913 |
| | Tmem164 | 542.956 | -0.220 | 0.061 | 10 | 0.045 | 1.652 |
| | Tmem196 | 166.953 | -0.713 | 0.188 | 7 | 0.029 | 2.044 |
| | Tmx2 | 1251.797 | -0.286 | 0.077 | 5 | 0.048 | 0.868 |
| | Tspan1 | 49.982 | 0.777 | 0.206 | 7 | 0.042 | 1.120 |
| | Ttbk2 | 69.348 | -1.574 | 0.349 | 6 | 0.009 | 1.120 |
| | Ttc39b | 71.332 | -0.981 | 0.256 | 5 | 0.027 | 1.959 |
| | Ttc9 | 521.426 | -0.276 | 0.073 | 6 | 0.035 | 1.652 |
| | Vcpip1 | 199.177 | -0.799 | 0.222 | 5 | 0.045 | 1.652 |
| | Wdr34 | 166.398 | 0.350 | 0.098 | 7 | 0.047 | 1.549 |
| | Xkr6 | 81.662 | 0.814 | 0.193 | 8 | 0.013 | 1.587 |
| | Zfp26 | 155.941 | -0.483 | 0.109 | 10 | 0.009 | 1.658 |
| | Zyx | 607.518 | 0.409 | 0.095 | 9 | 0.011 | 1.589 |

**Supplementary table 1.:** Parameters of significant RNA expression – behavior associations sorted by different sampling and analysis approaches. **Abbreviations:** log2FC: log2 fold-change, SE: standard error, FDR: false discovery rate, IHW: independent hypothesis weighting

| gene | robustness |
| --- | --- |
| Cytip | 3 |
| F7 | 3 |
| Hapln2 | 7 |
| Rragd | 9 |
| Scand1 | 9 |
| Zyx | 9 |
| Anp32a | 9 |
| Kit | 9 |
| Larp7 | 9 |
| Ccnjl | 9 |
| Gpx3 | 9 |
| Eral1 | 9 |
| Pdlim4 | 9 |
| Hmgn5 | 9 |
| Ccdc112 | 9 |
| Kcns2 | 9 |
| Slc25a10 | 9 |
| Taf3 | 9 |
| Adamts4 | 9 |
| Inpp5f | 10 |
| Pvrl2 | 10 |
| Tmem164 | 10 |
| Zfp26 | 10 |
| Cfdp1 | 10 |
| Orai1 | 10 |
| Tatdn3 | 10 |
| Cnih2 | 10 |

| | |
|---|---|
| Ppig | 10 |
| Pla2g7 | 10 |
| Krcc1 | 10 |
| Cdc42ep2 | 10 |
| Dgcr6 | 11 |
| Kcnj4 | 11 |
| Nat6 | 11 |
| Blcap | 12 |
| Samd4b | 12 |
| Col5a3 | 12 |
| Fam122a | 12 |
| Sec62 | 12 |
| Rnf145 | 13 |
| Abhd1 | 13 |
| Shbg | 13 |
| Eif3j | 13 |
| Sbno1 | 14 |
| Glb1 | 15 |

**Supplementary table 2. -** genes analyzed via qPCR in the amygdala, chosen for their robustness scores in the RNASeq analysis of mPFC tissue

| comparisons | gene symbol | r | p | Significance after FDR correction | aversive variables |
|---|---|---|---|---|---|
| PFC PCR vs RNASeq correlations | Adamts4 | 0.533578 | 4.15E-03 | * | - |
| | Col5a3 | 0.559829 | 2.39E-03 | * | - |
| | Glb1 | 0.608059 | 7.66E-04 | * | - |
| | Gpx3 | 0.580586 | 1.50E-03 | * | - |
| | Hapln2 | 0.741758 | 9.52E-06 | * | - |
| | Pdlim4 | 0.458861 | 1.61E-02 | * | - |
| | Pla2g7 | 0.568376 | 1.98E-03 | * | - |
| | Tatdn3 | 0.615385 | 6.34E-04 | * | - |
| AMY vs PFC expression level correlations | Glb1 | 0.647741 | 0.000259 | * | - |
| | Larp7 | 0.59655 | 0.001022 | * | - |
| | Pla2g7 | 0.574481 | 0.001725 | * | - |
| | Kcns2 | 0.482295 | 0.010843 | * | - |
| | Hmgn5b | 0.47779 | 0.011718 | * | - |
| | Tatdn3 | 0.473748 | 0.012553 | * | - |
| | Mrc1 | 0.407814 | 0.034722 | * | - |
| | Shbg | 0.397436 | 0.040087 | * | - |
| | Blcap | 0.40116 | 0.038091 | * | EPM_SuM |
| | Glb1 | 0.605006 | 0.000828 | * | LD_SiM |
| | Glb1 | 0.394383 | 0.041784 | * | COMP_SuM |

| | | | | | |
|---|---|---|---|---|---|
| AMY correlations with SiMs, SuMs, COMPs or aversive behavior | Inpp5f | -0.39178 | 0.043279 | * | EPM_aversive |
| | Pla2g7 | -0.58669 | 0.001297 | * | LD_SiM |
| | Pvrl2 | 0.388032 | 0.045499 | * | EPM_SiM |
| | Rragd | -0.39769 | 0.048988 | * | OF_aversive |
| | Shbg | 0.398459 | 0.048511 | * | OF_aversive |
| | Abhd1 | -0.4359 | 0.023036 | ns. | COMP_SuM |
| | Abhd1 | -0.40842 | 0.034426 | ns. | EPM_SuM |
| | Abhd1 | -0.40574 | 0.035746 | ns. | EPM_SiM |
| | Anp32a | 0.515873 | 0.005881 | ns. | EPM_SuM |
| | Ccnjl | -0.44375 | 0.020418 | ns. | EPM_aversive |
| | Cfdp1 | -0.42125 | 0.02865 | ns. | LD_SiM |
| | Cnih2 | 0.47619 | 0.012043 | ns. | EPM_SuM |
| | Cnih2 | 0.448176 | 0.019053 | ns. | EPM_SiM |
| | Glb1 | 0.47558 | 0.012169 | ns. | LD_SuM |
| | Gpx3 | 0.465201 | 0.014481 | ns. | EPM_SuM |
| | Gpx3 | 0.434066 | 0.023684 | ns. | LD_aversive |
| | Kit | -0.49112 | 0.009286 | ns. | EPM_aversive |
| | Larp7 | -0.42467 | 0.027248 | ns. | EPM_aversive |
| | Orai1 | -0.49611 | 0.008492 | ns. | EPM_SiM |
| | Orai1 | -0.44505 | 0.020007 | ns. | EPM_SuM |
| | Orai1 | 0.426129 | 0.026668 | ns. | LD_SiM |
| | Pla2g7 | -0.51709 | 0.005745 | ns. | LD_SuM |
| | Ppig | -0.43535 | 0.023227 | ns. | LD_SiM |
| | Rnf145 | -0.42467 | 0.027248 | ns. | EPM_aversive |
| | Samd4b | 0.470274 | 0.017678 | ns. | OF_aversive |
| | Samd4b | 0.384005 | 0.047986 | ns. | EPM_SuM |
| | Scand1 | -0.43224 | 0.024347 | ns. | EPM_aversive |
| | Tmem164 | 0.489391 | 0.009575 | ns. | EPM_SiM |
| | Tmem164 | 0.470696 | 0.013215 | ns. | EPM_SuM |
| | Tmem164 | 0.407814 | 0.034722 | ns. | LD_aversive |

| | | | | | |
|---|---|---|---|---|---|
| | Tmem164 | 0.400003 | 0.047566 | ns. | OF_aversive |
| | Zfp26 | -0.40296 | 0.037155 | ns. | EPM_aversive |
| | Zyx | 0.42735 | 0.02619 | ns. | LD_aversive |

**Supplementary table 3. -** validation of RNASeq experiments, correlations between amygdala expression and mPFC expression, or amygdala expression and anxiety
