## supplementary METHODS for "Improving anxiety research: novel approach to reveal trait anxiety through summary measures of multiple states"

### SUPPLEMENTARY MATERIALS AND METHODS

**Animals and housing.** Animals were kept in standard environmental conditions of 22±1°C temperature and 60±10% relative humidity in a 12:12h reversed circadian light-cycle room, with lights off at 8.00 a.m. Animals were housed in groups of 2-5 (mice and rats in separate rooms), rats in 1291H Eurostandard type III H cages (42.5x26.6x18.5 cm), and mice in 1284L EUROSTANDARD TYPE II L cages (36.5x20.7x14 cm). Water and food (Sniff, Soest, Germany) were available *ad libitum*. Experimental subjects underwent a 2-week acclimatisation period to their housing room, and also three 1-minute sessions of handling by the experimenter on three consecutive days right before experimentation. All experiments were approved by the Animal Welfare Committee of the Institute of Experimental Medicine, the National Scientific Ethical Committee on Animal Experimentation (permit no. PE/EA/00220-4/2022) and are in agreement with the 2010/63/EU directive.

**Behavioural tests.** All behavioural experiments were carried out in the first half of the animals' active phase by the same experimenter, in a testing room adjacent to the housing room. Animals were individually transferred from their homecage to the experimental room in a transfer-cage. Following the last behavioural test, animals were left undisturbed for 2 weeks, then swiftly terminated by decapitation in a non-stressed baseline state for later analysis of neurobiological correlates described below.

The Elevated Plus-Maze test consisted of an elevated plus-shaped testing arena with black plexiglass walls and a white plexiglass floor. The apparatus consisted of two open (aversive) and two enclosed (protected) arms connected by a central area (rat: height 70cm, arms 42x12 cm, centre 12x12 cm, closed arm wall height 35 cm; mouse: height 50cm, arms 30x7 cm, centre 7x7 cm, closed arm wall height 30 cm). Animals were placed in the centre of the testing arena facing the open arm in infrared lighting (13 lux), and were recorded for 5 minutes for later analysis. Aversive sampling was done in dim white lighting (49 lux).

The Light-Dark test consisted of an open, dimly lit (49 lux white light) arena with transparent plexiglass walls, and a closed, dark arena with black plexiglass walls and a black cover. The two chambers were connected by a narrow door (rat: light and dark boxes 50x50x40 cm; mouse: light and dark boxes 40x20x25 cm). Animals were placed in the light compartment and were recorded for 10 minutes. Aversive sampling was done in bright lighting (422 lux).

The Open Field test consisted of a large circular arena with black metal walls and a black wooden floor(for rats, diameter 100 cm, height ), or a white plastic box (for mice, 40x30x15 cm). The inner 50% (for mice) or 70% (for rats) of the arena was appointed as the centre. Animals were placed in the periphery of the arena in infra-red lighting, and were recorded for 10 minutes. Aversive sampling was done in the same experimental apparatus, but in dim white lighting (49 lux). The second type of Open Field testing (OF2) was done in a different experimental room by a different experimenter in dim white light. The testing apparatus was a black plastic square-shaped box (79x54x35 cm). Animals were placed in the periphery of the arena and recorded for 5 minutes.

The acoustic startle paradigm was performed with Long-Evans rats by a different experimenter 3 days after the last anxiety test. Startle reactivity was assessed in Plexiglas chambers (length: 25 cm, diameter: 12,5 cm) placed in sound-attenuating boxes (33 x 33x 48 cm) controlled by the SR-LAB software (SR-LAB Startle Response System, San Diego Instruments, USA) as previously described (Toth et al, 2014). Briefly, startle reactivity for increasing pulse intensity was assessed during a single session (approx. 20 min, light on in chambers). Session started with the delivery of 5 each of 120 dB startle pulses (over 65 dB background white noise) allowing startle to reach a stable level before specific testing. In a second block we presented four of each startle stimulus intensities (80, 90, 100, 110, and 120 dB) in a pseudorandom order with an average 15 sec intertrial intervals (range of 7-23 sec) between stimulus presentations. The average startle response for each intensity was calculated and considered as the index of startle reactivity.

The fear conditioning paradigm was carried out with Long-Evans rats 1 week after acoustic startle paradigm (Fig.4B). Fear conditioning was conducted in a different, brightly lit experimental room. Rats were placed in a 30x30x30 cm plexiglass chamber with metal grid floors. After a 2.5 min habituation period, 2.4 mA, 1 sec long, inescapable electric foot-shocks were delivered through the grid floor 10 times, each with a 30 sec inter-shock interval. Animals were placed in the same chamber 28 days later as a contextual reminder, then tested for fear-generalisation in a different apparatus with different contextual cues on the following day. Measured behavioural outcome was time spent freezing. Freezing was quantified by Ethovision as immobility lasting longer than 1 second, with detection thresholds and settings validated by significant correlation with expert hand-scoring ($r>0.9$).

For the social isolation protocol, upon weaning at P21, animals were reared in groups of 4 (social), or alone in a cage (isolated) for approx. 5 months until testing in adulthood.

**Statistical analysis of behaviour.** All statistical analysis was done in R statistical environment (version 3.6.2.)[39], through its integrated development environment, RStudio (version 1.3.1056)[40].

**Behavioural measures**:

*Light-Dark test scores as an example*

<u>calculation of scaled variables for aversive zone time (a.k.a. scaling):</u>

scaled(time in av. zone) = (individual's value – mean(population)) / SD(population)

<u>calculation of LD single measure z-scores (SiM) using variables from the first LD test:</u>

test1 SiM = scaled(time in av. zone) + scaled(entry frequency) - scaled(entry latency)

calculation of LD summary measure (SuM) using variables from all 3 LD tests:

SuM of 3 LD tests = average(test1 SiM, test2 SiM, test3 SiM)

calculation of LD-EPM-OF composite z-scores (COMP):

COMP = average(SuM(LD), SuM(EPM), SuM(OF))

**Repeatability analysis**. The analysis of test-repeatability was done for each test-type by calculating the proportion of within-test variance out of total variance[45], as shown below: *within test variance/(within test + between test variance),* where within-test (or between-individual) variance means the variation of the group's behaviour within a given test, and between-test (or within-individual) variance means variation of individuals' behaviour across weeks (test repetitions) for a given test-type. The R package 'rptR' (version 0.9.22)[46] was used to calculate variances and repeatability estimates similarly to previous investigations[47]. The 'rpt' function of the 'rptR' package relies on mixed-effects models that are fitted by functions of the R package 'lme4' (version 1.1.28)[48], and allows for estimation of raw variances and adjusted repeatability estimates. Inter-subject and inter-test variability were removed from the estimate by adding them to the model as fixed-effect covariates. The function also calculates 95% confidence intervals of estimates by parametric bootstrapping, with the number of parametric bootstraps for interval estimation set to 1000. Estimates with confidence intervals that did not include 0 were considered statistically significant. After examining normality of variables, the variances, adjusted repeatability estimates and their uncertainty were calculated for all behavioural variables (percent of time spent, entry frequency and latency into aversive zone, head-dipping, rearing).

**Random forest analysis.** To decrease the dimensions of our multivariate data and identify significant predictors of generalised fear in the CFP paradigm, we applied the semi-supervised machine learning method, Random Forest (RF)[49]. RF can rank the importance of each variable in the classification of experimental subjects to treatment groups. Since we model continuous behavioural readouts, instead of treatment groups, first we allocated subjects to high or low freezer categories. Our models produced 10000 decision trees and analyzed 4 features at a time to test which variable combination is the best to classify subjects to low or high freezing categories. Sampling was balanced to 10-10 subjects per group in the RF analysis. Variable importance was ranked based on permutation importance.

**Blood corticosterone analysis of Wistar rats.** Tail vein blood samples (0.3-0.5 ml) were collected in a resting state (5 days before the first experiment) and in stress-induced (immediately after OF2) conditions, and trunk blood was collected into ice-cold EDTA-containing tubes at the time of termination in a baseline state. After sampling, blood-containing tubes were centrifuged at 4 °C, and plasma was separated and stored at -20 °C until analysis. The quantification of plasma corticosterone was carried out using radioimmunoassay similarly to previous work by our laboratory [50]. Briefly, corticosterone was separated from corticosteroid-binding globulin (CBG) at low pH levels. CBG was kept inactive at low pH levels to avoid interference. Rabbit antiserum against corticosterone-3-carboxymethyloxime-bovine serum albumin was developed in the Institute, and 125I-labeled carboxymethyloxime–tyrosine–methyl ester was used as the tracer. All samples were measured in the same assay, with a sensitivity of 1 pmol/ml.

**RNA Sequencing of mPFC tissue.** Animals were swiftly decapitated after being transferred directly from their home-cage, their brains were removed, then cooled and washed in ice-cold saline. A 0.5 mm coronal section between Bregma 2.0 and 1.5 mm was sliced in a cooled slice matrix. Bilateral medial prefrontal cortex samples were dissected on ice as defined by the forceps minor of corpus callosum and the medial wall of the hemisphere (i.e. including the infralimbic, prelimbic and anterior cingulate cortices). Samples were immediately placed in Eppendorf tubes on dry ice and stored on -80C temperature until RNA isolation. Whole transcriptome-analysis via RNASeq was performed on the homogenised mPFC tissue. RNA sequencing was performed on 27 animals, chosen by their COMP scores. More precisely, a scale was created from all 54 animals' SuM scores, and every second animal in the scale was selected for RNASeq analysis. After the selection, correlations between the baseline composite score and behaviour in aversive conditions remained significant for both the selected and non-selected groups (Fig.5A). Analysis of gene-behaviour regressions was carried out using a negative binomial generalised linear model, with anxiety-scales generated from behaviour in different test-types (EPM/OF/LD/COMP) across test repetitions (SiM/SuM) as continuous covariates in the design. This was done to avoid losing relevant behavioural information by binarising the population into "high-anxiety" or "low-anxiety" groups. Percent of time spent with avoidance was used for SiMs, and composite scores of time and entry frequency were used for SuMs.

**Robustness.** We characterised gene-behaviour associations with the per unit of RNA read count change of an anxiety measure (SiM or SuM) and robustness. The robustness index is a measure to describe the fragility of a gene-behaviour association by giving the smallest number of randomly excludable samples that likely impair the significance of such a relationship. The index is calculated by a custom-written script that reanalyses the Spearman correlations between anxiety measures and RNA expression in randomly selected, different sample size sub-cohorts of the original population. We made 20000 correlations on all possible sample size sub-cohorts and calculated minimums, maximums, standard deviations and means of their p-values. We determined the cutoff criteria of a robustness index as the smallest sample size in which the standard deviation of the correlations' p-values reaches the 0.05 alpha level.

**Functional Clustering.** We applied the DAVID functional annotation tool that unites the 16 most frequently used annotation databases such as GO, KEGG or REACTOME.

Separate clusters consist of genes that share more similar functional labels in such databases. We showed the most inclusive functional clusters that possess significant enrichment of a function (Fig.7). Note that we excluded or merged functional categories that we considered too general or redundant, respectively (for details, see supplementary Table 1).

**qPCR analysis.** Total RNA was isolated using the RNeasy Lipid Tissue Mini Kit (Qiagen). The quality and quantity of the isolated RNA were assessed using the Qubit RNA BR Assay Kit and Qubit RNA IQ Assay Kit (Invitrogen, USA). A total of 500 ng of RNA was used for reverse transcription, which was performed using the High Capacity cDNA Reverse Transcription Kit (Thermo Fisher Scientific, USA). The concentrations of the resulting cDNA were determined using the Qubit ssDNA Assay Kit (Invitrogen, USA). Gene expression patterns were analyzed using custom 384-well TaqMan Gene Expression Array Cards (Applied Biosystems, USA). Each array card ran eight samples, including seven test samples and one run-control sample for calibration. The cDNA samples (200 ng each) were diluted and mixed with the TaqMan Gene Expression Master Mix to create a qPCR reaction mix, with a final concentration of 1 ng/µl. The qPCR reactions were performed using the ViiA 7 Real-Time PCR System (Applied Biosystems, USA), and the data were collected using the QuantStudio software (Applied Biosystems, USA).
